# irCLIP-RNP and Re-CLIP reveal patterns of dynamic protein associations on RNA

**DOI:** 10.1101/2024.09.27.615518

**Authors:** Luca Ducoli, Brian J. Zarnegar, Douglas F. Porter, Robin M. Meyers, Weili Miao, Nicholas M. Riley, Suhas Srinivasan, Leandra V. Jackrazi, Yen-Yu Yang, Zhouxian Li, Yinsheng Wang, Carolyn R. Bertozzi, Ryan A. Flynn, Paul A. Khavari

## Abstract

RNA binding proteins (<u>RBPs</u>) control varied processes, including RNA splicing, stability, transport, and translation^1-3^. Dysfunctional RNA-RBP interactions contribute to the pathogenesis of human disease^1,4,5^, however, characterizing the nature and dynamics of multiprotein assemblies on RNA has been challenging. To address this, non-isotopic ligation-based ultraviolet crosslinking immunoprecipitation^6^ was combined with mass spectrometry (<u>irCLIP-RNP</u>) to identify RNA-dependent associated proteins (<u>RDAPs</u>) co-bound to RNA with any RBP of interest. irCLIP-RNP defined landscapes of multimeric protein assemblies on RNA, uncovering previously unknown patterns of RBP-RNA associations, including cell-type-selective combinatorial relationships between RDAPs and primary RBPs. irCLIP-RNP also defined dynamic RDAP remodeling in response to epidermal growth factor (<u>EGF</u>), uncovering EGF-induced recruitment of UPF1 adjacent to HNRNPC to effect splicing surveillance of cell proliferation mRNAs. To identify the RNAs simultaneously co-bound by multiple studied RBPs, a sequential immunoprecipitation irCLIP (<u>Re-CLIP</u>) method was also developed. Re-CLIP confirmed binding relationships seen in irCLIP-RNP and detected simultaneous HNRNPC and UPF1 co-binding on RND3 and DDX3X mRNAs. irCLIP-RNP and Re-CLIP provide a framework to identify and characterize dynamic RNA-protein assemblies in living cells.

---

Ultraviolet light-induced crosslinking (**<u>UVC</u>**) followed by RBP immunoprecipitation and RNA-sequencing (**<u>CLIP-seq</u>**) captures direct RBP-RNA interactions^7,8^ and can localize RNA binding sites for RBPs studied^9^ but is not designed to study the proximal proteins assembled on shared RNA targets. Relationships among the diverse repertoire of RBPs that assemble on their target RNAs thus remain poorly characterized, highlighting the need for new approaches to characterize multiprotein assemblies on RNA. Two methods were designed to address this. **<u>irCLIP-RNP</u>** uses non-isotopic infrared-CLIP (**<u>irCLIP</u>**)^6^ with biotin/infrared dye adapters for 3’ RNA ligation, along with lower concentrations of RNase for in-lysate digestion, to visualize RNA-dependent *cis*-RBP assemblies and identify their protein components with mass spectrometry (**<u>MS</u>**). **<u>Re-CLIP</u>** employs two independent rounds of irCLIP immunoprecipitation followed by RNA sequencing to identify the specific RNA molecules simultaneously bound by multiple RBPs of interest. These methods are designed to facilitate the study of the composition, functions, and dynamics of combinatorial protein assemblies on RNA.

## Complexity of protein associations on RNA

To understand if CLIP-based methods could uncover evidence of multiprotein assembly of proteins on shared target RNAs, a panel of well-studied RBPs involved in RNA production, maturation, and function was examined by irCLIP. RNase A was titrated across a range of concentrations to assess RNA dependency of larger molecular weight (**<u>MW</u>**) RNA-protein complexes. Infrared images of nitrocellulose-transferred irCLIP-ligations displayed signal above the expected MW of the target RBP (“monomeric RBP zone”) for all RBPs, even at the highest RNase concentrations (**Fig. 1a, Extended Data Fig. 1a**). This is expected as the target RBP is covalently crosslinked to an RNase A-trimmed piece of RNA and ligated to the irCLIP adaptor, shifting the RNP above the endogenous MW of the RBP. This was further supported by immunoblotting the HNRNPC and TARDBP RBPs, where RNase titration generated multiple bands that were shifted to a higher MW (**Extended Data Fig. 1b**), suggesting that each protein was captured in different-sized complexes. Quantifying the total amount of irCLIP signal at each dose of RNase A normalized per cell unit demonstrated that an intermediate amount of RNase A, 55ng/mL, generated the maximal amount of successful ligation events for HNRNPC, HNRNPA2B1, and FUS (**Extended Data Fig. 1c**). Distribution of the irCLIP ligations across 16 RBPs studied by irCLIP showed that intermediate RNase treatment resulted in a large fraction of these ligations occurring outside the monomeric RNP zone (**Fig. 1b**), suggesting capture of diverse RBP-RNA associations dependent on the RBP of interest.

**Figure 1:**
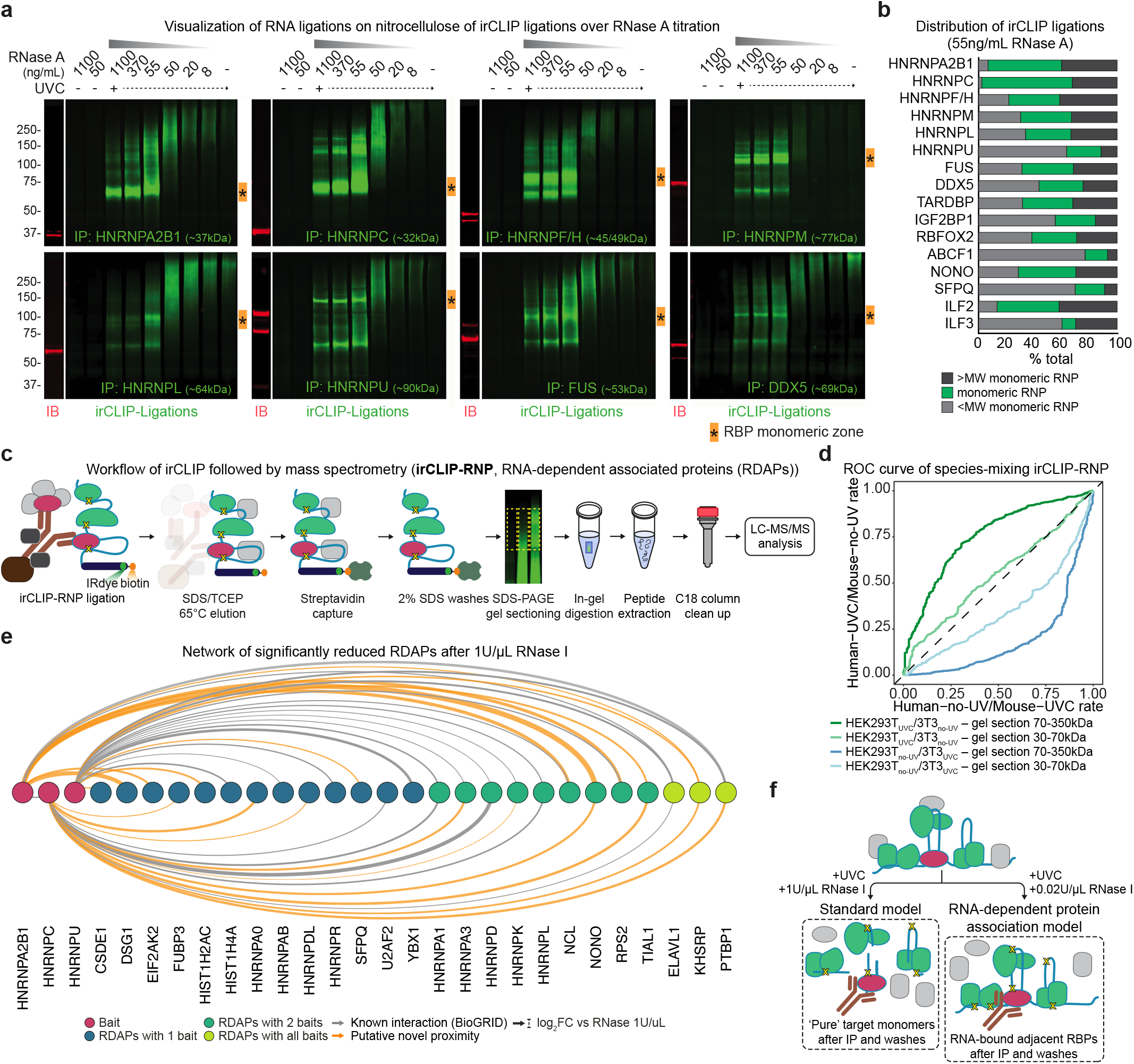
Complexity of protein associations on RNA. **(a)** Nitrocellulose images indicate the infrared signal of irCLIP ligations subsequent to in-lysate digestion with six RNase A concentrations (from 1100 ng/mL to 8 ng/mL) in HEK293T cells for different RBPs involved in RNA processing. IP: immunoprecipitation; UVC: UV crosslinking; green signal: irCLIP ligation; red signal: immunoblotting for the tested RBP; asterisks in orange box: RBP monomeric zone. The reported kDa for each RBP represents the calculated MW. **(b)** Stacked bars indicate the distribution of irCLIP ligation infrared signal between the monomeric RNP gel section (green) and the regions with higher (black) or lower (grey) MW (MW) after in-lysate digestion with 55ng/mL RNase A. irCLIP ligations were analyzed in the gel region between 60kDa and 350kDa. **(c)** Workflow to perform irCLIP followed by mass spectrometry (**<u>irCLIP- RNP</u>**) to uncover RNA-dependent associated proteins (RDAPs). **(d)** Lines indicate ROC curves of cell-type-specific unique peptide intensities after species-mixing irCLIP-RNP for HNRNPC (UVC human lysis; no-UV mouse lysis) and HNRNPA2B1 (no-UV human lysis; UVC mouse lysis). Green and light-green lines: high and low gel section in HNRNPC irCLIP-RNP. Blue and light blue lines: high and low gel section in HNRNPA2B1 irCLIP-RNP. **(e)** Arc plot of HNRNPC, HNRNPA2B1, and HNRNPU (dark fuchsia nodes) and the 25 RDAPs that are significantly reduced after 1U/µL RNase I (dark blue, green, and light-green nodes). Grey edges: known interactions according to BioGRID database; orange edges: novel proximity associations; edges width: log_2_FC value versus RNase 1U/µL. **(f)** Schematic representation of complex RNA-dependent protein association after irCLIP-RNP purification using intermediate RNase I treatment.

While most RBPs investigated displayed subtly different profiles, irCLIP patterns of the cognate heterodimeric pairs, SFPQ:NONO^10^ and ILF2:ILF3^11^, generated similar RNase digestion patterns, suggesting that both proteins contribute to each other’s CLIP ligations (**Extended Data Fig. 1a**). Consistent with this, protein co-immunoprecipitation was observed under standard ENCODE CLIP conditions not only for SFPQ:NONO and ILF2:ILF3 heterodimers, but also among members of the large assembly of splicing regulators (**<u>LASR</u>**) complex (HNRNPM, HNRNPC, and ILF2:ILF3)^12^ (**Extended Data Fig. 1d**). While these pairs of RBPs represent strong known heterodimers, they raise the possibility that, in some cases, proteins present in the higher MW region of the gel under intermediate RNase conditions would be reduced upon higher RNase treatments if their slow migration in the gel was due to RNA-association.

To test this, irCLIP^6^ was coupled with in-gel peptide isolation followed by mass spectrometry, (**<u>irCLIP-RNP</u>**) (**Fig. 1c**). irCLIP-RNP uses UVC crosslinking, intermediate RNase I digestion, and stringent washes to detect RNA-dependent associated proteins (**<u>RDAPs</u>**) that are covalently bound to RNA with the primary immunoprecipitated RBP, with RNP visualization enabled by infrared 3’ RNA labeling (adapter length: 78nt – approx. 25kDa). To evaluate the method’s reproducibility, we first performed irCLIP-RNP of HNRNPC by comparing UVC and non-UV treated (**<u>no-UV</u>**) samples from two gel sections ranging from 30-60kDa, which we termed “free RNA ligation zone”, and 60-350kDa, termed “whole RNP zone”, as it covers the entire region of irCLIP-RNP signal (**Extended Data Fig. 1e**). While RNase A and RNase I yielded similar digestion profiles, RNase I was selected in irCLIP-RNP for its reproducible digestion patterns at intermediate concentrations. Infrared in-gel scan showed consistent signals across UVC as well as no-UV samples (**Extended Data Fig. 1f**). Correlation analysis of the 140 detected proteins displayed a high correlation between UVC samples as well as between no-UV samples coming from the whole RNP subzone (**Extended Data Fig. 1g**). Differential enrichment analysis identified 45 proteins to be enriched in the UVC whole RNP zone, nominating them as bona fide RDAPs (**Extended Data Fig. 1h, Supplementary Table 1**). Several HNRNP family members were identified, as well as other known RBPs, such as KHSRP, ELAVL1, TARDBP, NONO, and SFPQ. Next, non-specific RNA binding during cell lysis was assessed by performing irCLIP-RNP of free RNA ligation and whole RNP zones after mixing lysates from human (HEK293T) and mouse (3T3) cells. UVC-human samples were compared to no-UV mouse and vice versa (**Extended Data Fig. 1i-j**). Receiver operating characteristic (**<u>ROC</u>**) analysis of cell-type-specific unique peptide intensities in UVC human/no-UV mouse and no-UV human/UVC mouse samples showed that irCLIP-RNP selectively enriched for peptides from the respective crosslinked cell type in the highest RNP subzone, with minimal cross-species contamination (**Fig. 1d**). More importantly, this enrichment was more prominent in the whole RNP zone, further strengthening the capability of irCLIP-RNP to identify crosslink-dependent RDAPs. This pattern was evident after analysis of the distribution of unique peptides across human and mouse protein sequences (**Extended Data Fig. 1k**).

The composition of the whole RNP zones between two RNase I concentrations (1U/µL and 0.02U/µL) was next compared in HEK293T cells for three members of well-characterized heterogeneous nuclear ribonuclear protein (**<u>hnRNP</u>**) complexes (HNRNPA2B1, HNRNPC, and HNRNPU^13-15^) (**Extended Data Fig. 2a**). Visualization of irCLIP-RNP ligations on nitrocellulose membranes showed distinct MW patterns between high (1U/µL) and intermediate (0.02U/µL) RNase I treatments (**Extended Data Fig. 2b**). The whole RNP zones were subjected to in-gel digestion followed by MS. irCLIP-RNP detected 33, 43, and 53 UVC-enriched proteins for HNRNPA2B1, HNRNPC, and HNRNPU, respectively (**Extended Data Fig. 2c, Supplementary Table 2**). Differential enrichment analysis between high (1U/µL) and intermediate (0.02U/µL) RNase I samples identified 27 unique proteins that displayed reduced protein intensity after high RNase I treatment in at least one of the tested RBPs. Specifically, 6, 15, and 20 proteins were significantly reduced upon high RNase I treatment in either HNRNPA2B1, HNRNPC, and HNRNPU samples, revealing different patterns of RBP co-associations, where some factors retained or lost their associations after high RNase I treatment, depending on the immunoprecipitated RBP (**Extended Data Fig. 2d-e**). Although substantial overlap of RDAPs was observed among hnRNP proteins studied, each hnRNP also displayed distinct sets of RDAPs reduced after 1U/µL RNase I digestion (**Fig. 1e, Extended Data Fig. 2f**). Some of the proteins identified were already known to interact with the tested RBPs^16^, supporting the validity of the detection approach. irCLIP-RNP also identified many RDAPs showing putative novel RNA-dependent proximity with the primary hnRNP RBPs (**Fig. 1e**), highlighting the capacity of irCLIP-RNP to decipher patterns of protein associations on RNA.

**Figure 2:**
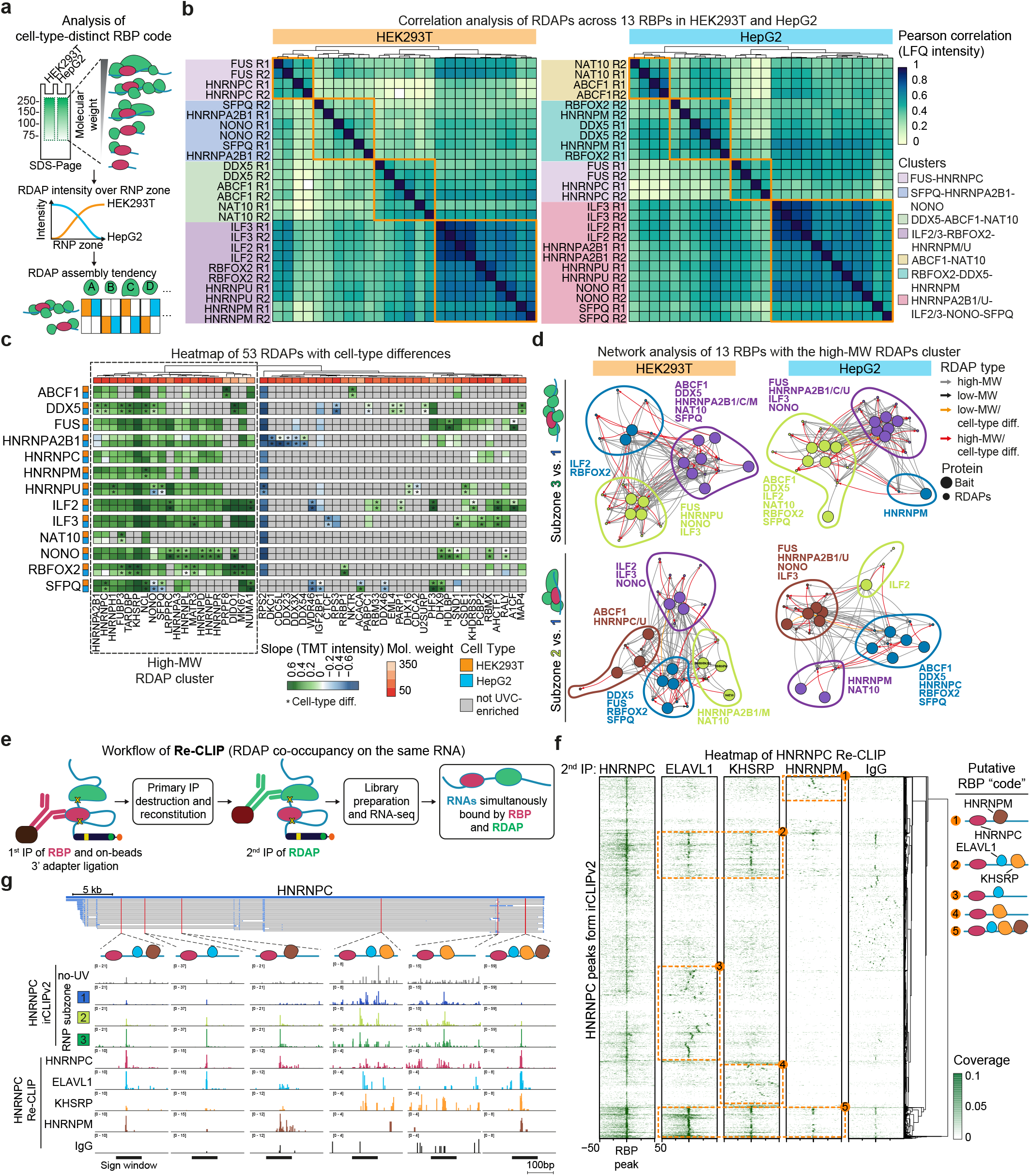
irCLIP-RNP reveals cell-type-selective RDAP patterns and co-binding of RNAs. **(a)** Workflow of the characterization of RDAP assembly tendency between HEK293T and HepG2 by analyzing the distribution of irCLIP-RNP signal across the whole RNP zone. **(b)** Heatmap indicates the Pearson correlations calculated from imputed label-free quantitation (**<u>LFQ</u>**) intensities for the 360 UVC-enriched RDAPs in 13 RBPs (HEK293T: left; HepG2: right). Clustering was performed using correlation distance and Ward clustering methods. **(c)** Heatmap indicates slope values calculated from centered TMT intensities from the RNP subzones #1-3 for the 53 RDAPs showing significant differences in irCLIP-RNP signal distribution between HEK293T (orange) and HepG2 (blue). ^*^: cell-type-selective RDAP with FDR < 0.05 in the corresponding RBP as determined by differential enrichment analysis. Clustering was performed using Euclidean distance and Ward clustering methods, identifying 4 clusters (orange boxes). **(d)** Network schematics of high-MW proteins. Community detection was performed after applying a Louvain clustering algorithm. Grey edge: high-MW protein; black edge: low-MW protein; orange edge: cell-type differential low-MW proteins; red edge: cell-type differential high-MW proteins. Dot size represents bait or RDAPs. **(e)** Workflow of Re-CLIP approach for the detection of co-occupancy of RDAPs on the same RNA molecules. After the first immunoprecipitation and 3’ ligation, RNP ligations are subjected to secondary immunoprecipitation followed by library preparation and RNA sequencing. **(f)** Heatmap indicates normalized coverage around RBP peaks of Re-CLIP. Orange boxes highlight regions that are co-bound by HNRNPC and RDAPs. Clustering was performed using Euclidean distance and Ward clustering methods **(g)** IGV tracks indicate normalized coverage of Re-CLIP samples at several binding sites across the HNRNPC transcript. Tracks show the signal sum of 2 replicates.

The loss of a subset of putative RDAPs at higher RNase concentrations suggested that RDAPs may reside at differing nucleotide distances on co-bound RNA molecules from pulled-down RBPs (**Fig. 1f**). Consistent with this, analysis of RNA size resulting from HNRNPC irCLIP-RNP gel sections ranging from 60-120kDa, 120-225kDa, and 225-350kDa (referred to as RNP subzones #1-3) showed increasing RNA molecule length ranging from 10 to ∼210 nucleotides with increasing RNP MW (**Extended Data Fig. 2g**). The MW shift observed after intermediate RNase I treatment, however, was not due only to the change in RNA length (∼15-140nts, ranging from ∼5-40 kDa) but is likely also due in part to protein complexes crosslinked to the same RNA as the immunoprecipitated RBPs (**Extended Data Fig. 2g-h**). Intermediate RNase treatment, therefore, generates distinct MW patterns that represent a heterogeneous collection of RDAPs associated with the primary immunoprecipitated RBP on the same RNA molecules.

## irCLIP-RNP reveals cell-type-selective RDAP patterns for specific RBPs

The MW shift observed after intermediate RNase suggested a distribution of different kinds of RDAP complexes. RDAP distribution over the regions of irCLIP-RNP signal was accordingly examined to assess the tendency of RDAPs to occur either in low or high MW complexes in different cell contexts (**Fig. 2a**). Label-free irCLIP-RNP was first performed on entire gel sections (“whole RNP zone”; **Extended Data Fig. 3a**) in two cell lines (HEK293T and HepG2) for 13 RBPs involved in RNA processing and stability. This identified 360 total UVC-dependent RDAPs in at least one cell line for the RBPs tested (**Extended Data Fig. 3b-c, Supplementary Table 3**). Regression analysis of previously published protein expression data^17^ showed comparable expression levels for these RDAPs between HEK293T and HepG2 cells (**Extended Data Fig. 3d**), indicating the different protein abundance does not account for observed differences. Gene ontology (GO) analysis (**Extended Data Fig. 3e**) showed that a large fraction of these proteins displayed RNA-binding functions. Correlation analysis of the 360 RDAPs identified several RBP clusters, with composition differences between the two cell lines (**Fig. 2b**). For instance, HNRNPA2B1, NONO, SFPQ, and RBFOX2 followed a more HNRNPU/ILF2/3-like interactome, depending on the cell studied (**Fig. 2b**). This was further supported by reciprocal pull-down between the 13 RBPs, which showed cell-type context dependency (**Extended Data Fig. 3f**). In fact, NONO and HNRNPU showed higher reciprocal pull-down efficiencies in HepG2 cells, which resulted in higher interactome correlation compared to HEK293T (**Fig. 2b, Extended Data Fig. 3f**). These data indicate that patterns of RBP-RDAP binding relationships on RNA may be influenced by cellular context.

**Figure 3:**
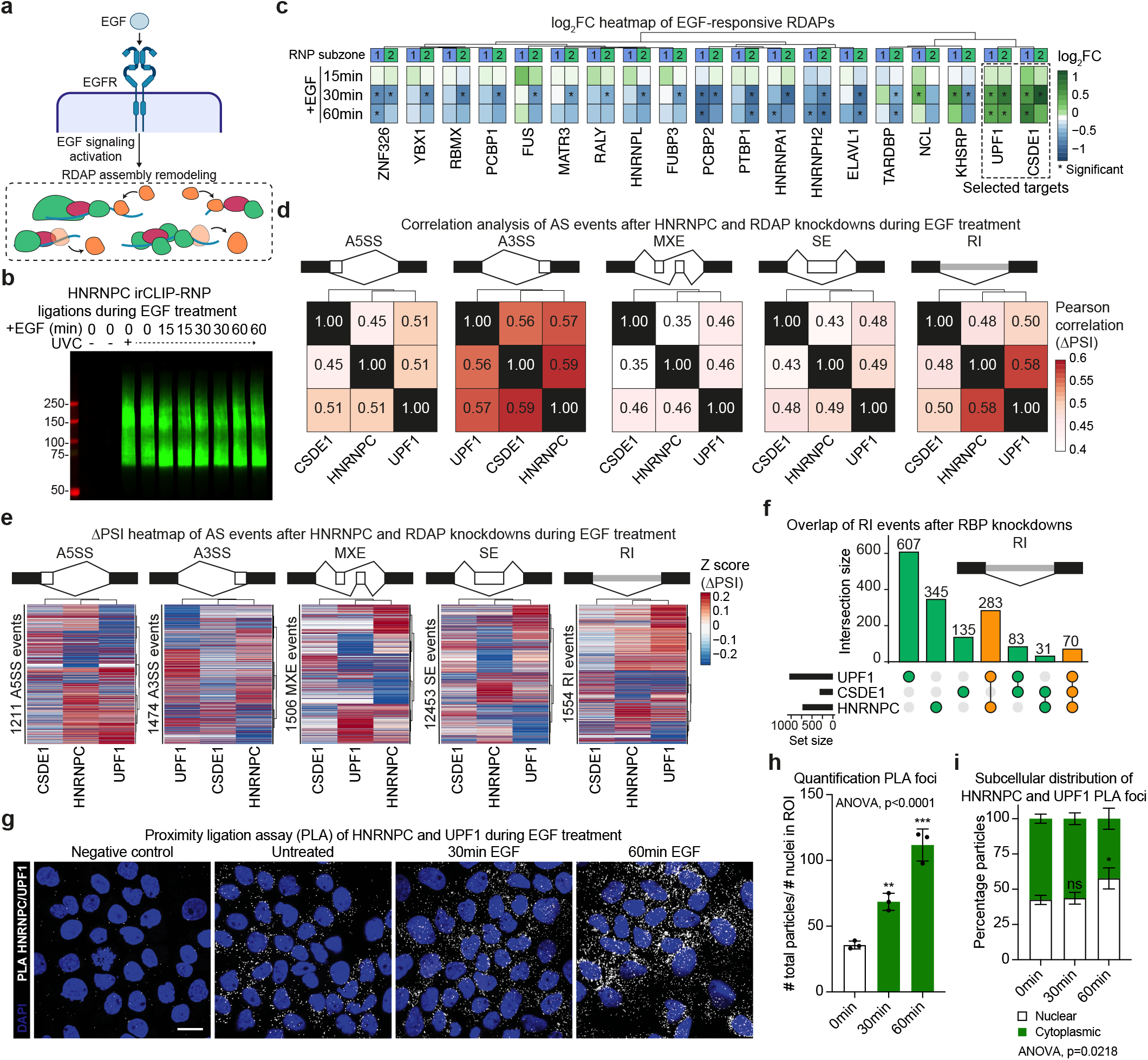
irCLIP-RNP reveals dynamic remodeling of RDAPs. **(a)** Schematic representation of EGF signaling activation effects on the remodeling of the RBP-related RDAP landscape. **(b)** Nitrocellulose image indicates HNRNPC irCLIP-RNP ligation signal of EGF-timecourse samples (n = 2). **(c)** Heatmap indicates log_2_FC values against time point 0 calculated from TMT intensities in RNP subzones #1-2 for the 19 RDAPs showing significant perturbation over EGF stimulation. ^*^: EGF-responsive RDAP with FDR < 0.05 and an abs(log_2_FC) > 0.3 in the related time point. Clusters were determined using Mahalanobis distance and Ward clustering methods. **(d)** Heatmaps indicate Pearson correlation coefficients calculated from ΔPSI values of significant AS events (A5SS, A3SS, MXE, RI, and SE) in EGF-stimulated samples after HNRNPC and RDAPs knockdowns. Correlation was calculated on significant AS events in at least one knockdown sample compared to control siRNAs. Clustering was performed using correlation distance and “complete” clustering algorithm. **(e)** Heatmaps indicate ΔPSI values of significant AS events in EGF-stimulated samples after HNRNPC and RDAPs knockdowns. Clustering was performed using correlation distance and Ward (rows) and “complete” (columns) clustering methods. **(f)** Bars indicate the overlap of significant RI events between HNRNPC and RDAP knockdown samples after EGF stimulation. **(g)** Representative images of PLA signal for HNRNPC and UPF1 in untreated and EGF-treated (30 and 60min) A431 cells. DAPI (blue signal) was used to outline the nuclear shape. Scale bars represent 20 µm. **(h)** Bars indicate the PLA foci per cell of HNRNPC and UPF1 in untreated and EGF-treated (30 and 60min) A431 cells. **(i)** Stacked bars indicate the nuclear and cytoplasmic PLA signal distribution in EGF-treated (0, 30, and 60min) A431 cells. In **h** and **i**, data are mean ± s.d; ^*^P-value < 0.05, ^**^P-value < 0.01, ^***^P-value < 0.001, ns: not significant – one-way ANOVA with Tukey’s test against untreated cells (n = 3).

In parallel, to examine RDAPs by MW zones, irCLIP-RNP was performed, followed by multiplexed isobaric tandem mass tag (**<u>TMT</u>**) quantitative MS in HEK293T and HepG2 cells for the 13 RBPs with the three RNP subzones #1-3 analyzed separately (**Extended Data Fig. 3g**). Of the 360 UVC-enriched proteins identified above, 163 RDAPs were also enriched in the irCLIP-RNP TMT dataset (**Supplementary Table 4**), supporting their association with the primary RBPs studied. Differential enrichment analysis between the three RNP subzones identified many RDAPs showing significantly higher TMT intensities in either low-MW or high-MW gel sections in the two cell types (**Extended Data Fig. 4a**). Consistent with the MW model, RDAPs categorized as “low” or “high” displayed significantly lower or higher MW compared to background proteins (**Extended Data Figure 4b**). However, these large MW proteins were still well below the upper limit of 350kDa gel sectioning (**Extended Data Figure 4b**). Thus, the shift seen at the higher sections in the gel represents a heterogenous mixture of RNA-bound complexes likely composed of large and/or small proteins, as schematized in **Fig. 2a**. Among these, 53 RDAPs showed a cell-type-dependent distribution across the gel lane where irCLIP-RNP signal was present in at least one of the RBPs tested (**Extended Data Fig. 4a**).

**Figure 4:**
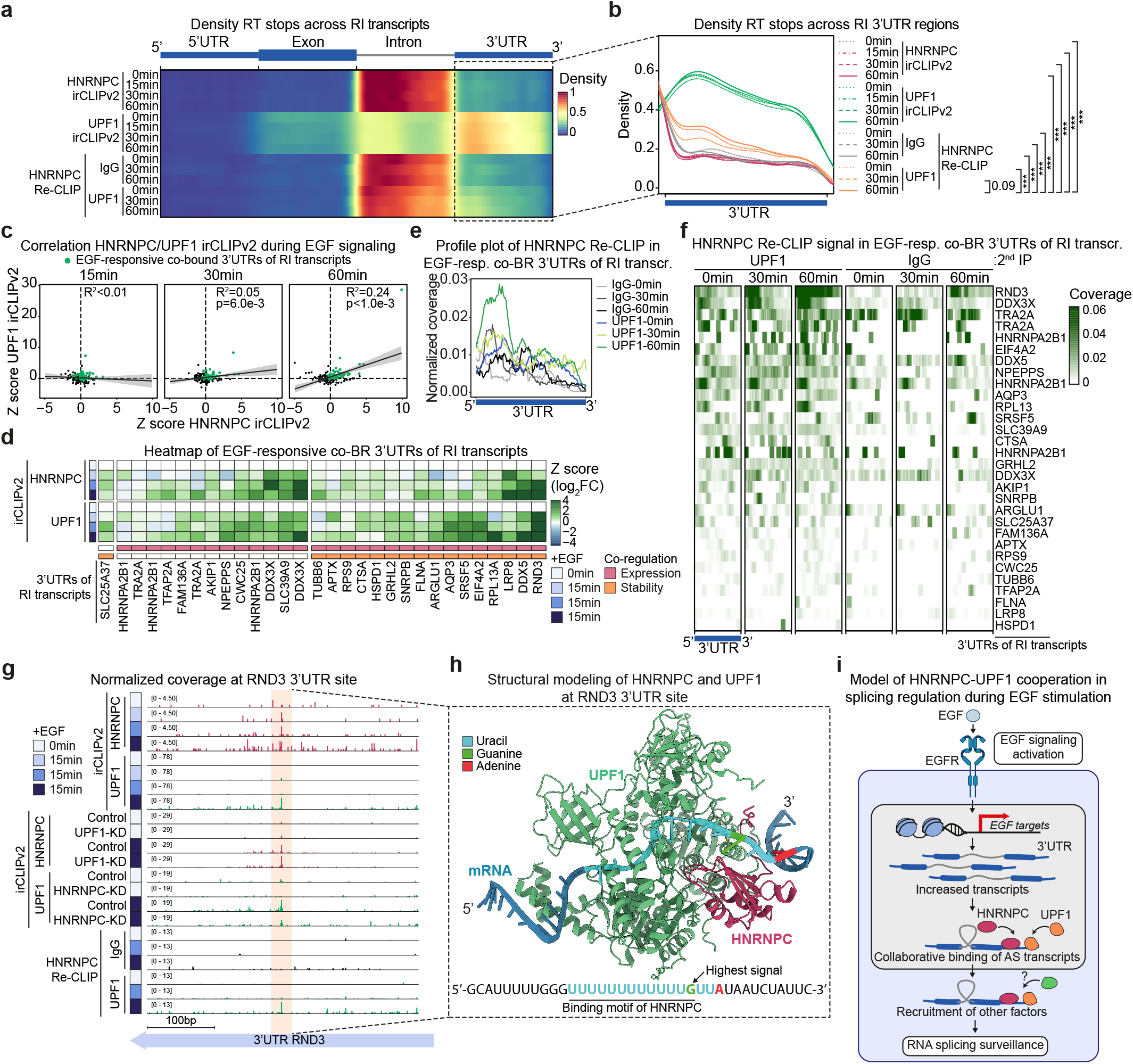
UPF1 supports HNRNPC RNA splicing surveillance of EGF-regulated mRNAs. **(a)** Heatmap indicates the density of RT stops across RI transcript regions (5’UTR, exon, intron, and 3’UTR) of HNRNPC and UPF1 irCLIPv2 and HNRNPC Re-CLIP of UPF1 data. **(b)** Lines indicate RT stop density in 3’UTR regions magnified from **a**. ^***^P-value < 0.001 – the Kolmogorov-Smirnov test between 60min-EGF-treated HNRNPC-UPF1 re-CLIP samples and displayed comparisons. **(c)** Scatterplot indicates Z score values estimated against time point 0 from total intensity across the 3’UTR regions of RI transcripts in HNRNPC and UPF1 irCLIPv2 data. Green dots: co-bound 3’UTRs of RI transcripts having a max(log_2_FC) > 0 across EGF stimulation in both irCLIPv2 datasets. **(d)** Heatmap indicates Z score values across EGF-stimulated HNRNPC and UPF1 irCLIPv2 samples of the 30 EGF-responsive 3’UTRs of RI transcripts that are co-bound and co-regulated (co-BR) by HNRNPC and UPF1. Shades of blue boxes: EGF stimulation time points; pink box: co-regulated expression; orange box: co-regulated stability. **(e)** Lines indicate normalized coverage of HNRNPC-UPF1 Re-CLIP during EGF stimulation across EGF-responsive co-BR 3’UTRs of RI transcripts. Coverage on the negative strand was reversed to be represented in a 5’ to 3’ orientation. Shades of grey: HNRNPC-IgG Re-CLIP; blue/green/light-green lines: HNRNPC-UPF1 Re-CLIP. **(f)** Heatmap indicates normalized coverage of HNRNPC-UPF1 Re-CLIP across EGF-responsive co-BR 3’UTRs of RI transcripts. **(g)** IGV tracks indicate normalized coverage of HNRNPC and UPF1 irCLIPv2 time-course/knockdown and HNRNPC-UPF1 Re-CLIP samples at 3’UTR of RND3. Tracks show the signal sum of 2 replicates. Shades of blue boxes: EGF stimulation time points. **(h)** *De novo* structure prediction using AlphaFold 3 and manual refinement provides a three-dimensional model of the protein-RNA multimer complex comprising of HNRNPC, UPF1, and 3’ UTR mRNA of RND3. **(i)** Model for the collaboration between HNRNPC and UPF1 in RNA splicing surveillance during EGF signaling.

The distribution of the 53 identified RDAPs was then analyzed by calculating the slope from TMT intensities across the three RNP subzones (**Extended Data Fig. 4c**) where RDAPs occurring in the higher section of the irCLIP-RNP signal distribution display a higher slope value than RDAPs at a lower section. Hierarchical clustering of these values further supported the context dependency of these associations, especially for DDX5, HNRNPU, NONO, SFPQ, and RBFOX2 (**Fig. 2c**). More importantly, this analysis highlighted the distribution changes between HEK293T and HepG2 cells of a consistent cluster of 20 RDAPs occurring at higher gel sections. This cluster comprised several hnRNPs, including components of the LASR complex (HNRNPM, HNRNPH1, HNRNPC, MATR3)^12^ as well as other well-known RBPs, such as NONO, SFPQ, TARDBP, KHSRP, and ELAVL1, which displayed distinct cell-type-related modular distribution according to the primary RBP immunoprecipitated (**Fig. 2c**). In particular, HNRNPU showed a clear difference in the distribution for NONO and SFPQ between HEK293T and HepG2, further highlighting the cell-type context dependency of the association between these factors (**Fig. 2c, Extended Data Fig. 3h**). This was also seen after network analysis followed by community clustering of the 13 RBPs with the members of the high-MW RDAP cluster which showed distinct patterns between cells lines as well as between gel sections (**Fig. 2d**). Nitrocellulose images of sequential irCLIP immunoprecipitation of NONO and some of its RDAPs, along with negative controls THOC2 and EIF5B, showed signal for SFPQ, HNRNPC, HNRNPU, and HNRNPM (**Extended Data Fig. 4d**), consistent with the premise that these associations occur on the same individual RNA molecules.

To examine the RBP-RDAP proximity on co-bound RNA, we performed a refined version of the original irCLIP method^6^, termed **<u>irCLIPv2</u>**, by introducing in-solution capture of nitrocellulose liberated ligated RNPs, solid-support cDNA synthesis, and 3’ cDNA-tailing, which reduced the library construction time to less than 8 hours (**Extended Data Fig. 5a**). irCLIPv2 ligations from HEK293T cells underwent library preparation then RNA sequencing for each of the three RNP subzones (#1-3) for the RDAPs KHSRP, ELAVL1, and HNRNPFH, and 4 members the HNRNPs protein family (**Extended Data Fig. 5a-b**). KHSRP and ELAVL1 were studied first because of their previously shown tendency to associate with RNA in multiprotein complexes^18,19^. irCLIPv2 of the ALYREF protein was included as a negative control since no selected RBPs showed an association with this protein in irCLIP-RNP data. After mapping and crosslink site counting, the DEWSeq differential sliding window approach was used to select significantly enriched regions in the irCLIPv2 samples against no-UV control (**Extended Data Fig. 5a, Supplementary Table 5**). KHSRP and ELAVL1 showed the highest percentage of the 144,879 total binding regions and the broadest binding densities of RNA targets, further supporting extensive binding across the transcriptome by these RBPs (**Extended Data Fig. 5c-d**). Moreover, significant overlap was found between irCLIPv2-identified regions and previously reported easyCLIP sites for HNRNPC and RBFOX2^20^ (**Extended Data Fig. 5e**). Analysis of binding profiles at the highest crosslink site of the RBPs and RDAPs showed a general 3’-shift of the signal in the higher RNP subzones 2 and 3, suggesting that this is potentially due to RDAP co-binding on the same RNA molecule, resulting in early truncation during library preparation (**Extended Data Fig. 5f-g**).

Next, each region was categorized according to its slope values in the three RNP subzones into final low, medium, or high binding regions (**Extended Data Fig. 5h**). As expected, RBPs displayed distinct binding patterns across known functional RNA sequences (**Extended Data Fig. 5i**). For instance, the intron-binding preferences for HNRNPC, HNRNPM, and KHSRP and the 3’UTR binding preferences for ELAVL1 were consistent with previous data^21^ with higher levels in the regions categorized as “high” (**Extended Data Fig. 5i**). Guided by RNA sizing results (**Extended Data Fig. 2g-h**), motif enrichment analysis was performed of low, medium, and high binding regions, which revealed that irCLIPv2 retrieved RBP binding motifs of most high-MW proteins, with congruent enrichment pattern across RNP subzones as seen in irCLIP-RNP data (**Extended Data Fig. 5j-k**), further indicating that irCLIP-RNP detects RNA-dependent proteins assembled on co-bound RNAs. Indeed, analysis of the nearest binding regions between HNRNPC and RDAPs showed higher log_2_FC against no-UV values at HNRNPC peaks in the RNP subzones #3 near the RDAPs (**Extended Data Fig. 5l**), supporting the observation that those RDAPs, especially KHSRP and ELAVL1, are mainly associated with HNRNPC in the highest RNP subzone and, therefore, may co-bind the same RNA molecules. Consistent with this premise, RNA pull-down of in-vitro transcribed RNA of KHSRP-HNRNPC regions previously incubated with HEK293T total cell lysate showed enrichment of both proteins at two different HNRNPC-KHSRP binding pairs (**Extended Data Fig. 5m-n**).

To provide a higher throughput way of identifying RNA molecules that are simultaneously co-bound by multiple RBPs of interest, irCLIPv2 was adapted to perform sequential immunoprecipitation followed by RNA sequencing (**<u>Re-CLIP</u>**) (**Fig 2e**). After RNase digestion as well as the first immunoprecipitation and 3’ ligation, the resulting pull-down material underwent a second immunoprecipitation, with Re-CLIP ligations then subjected to RNA sequencing (**Fig 2e**). To explore RNA-dependent RBP-RDAP multifactor associations, Re-CLIP was performed in HEK293T cells. HNRNPC was first immunoprecipitated followed by separate second immunoprecipitations for ELAVL1, KHSRP, and HNRNPM (**Extended Data Fig. 5o**). Nitrocellulose imaging showed efficient re-immunoprecipitation of HNRNPC itself as well as consistent signal for ELAVL1 and KHSRP re-immunoprecipitation (**Extended Data Fig. 5p**). Clustering analysis using HNRNPC binding sites identified in the primary irCLIPv2 dataset showed coherence of re-immunoprecipitated HNRNPC with HNRNPC primary irCLIPv2 (**Fig. 2f**). Of interest, multiple subsets of HNRNPC binding regions showed different modularity with the RDAPs, suggesting that Re-CLIP uncovered discrete combinatorial RNA binding relationships between HNRNPC and its specific RDAPs (**Fig. 2f**). For example, the HNRNPC RNA, which is a well-known binding target of the HNRPNC protein itself, showed diverse combinations of HNRNPC co-binding with ELAVL1, KHSRP, and HNRNPM at different regions along the HNRNPC RNA transcript (**Fig. 2g**). Thus, by localizing RBPs and RDAPs on the same RNA molecules that they co-bind, Re-CLIP may further extend the understanding of RBP-RDAP co-binding relationships identified by irCLIP-RNP.

## irCLIP-RNP reveals dynamic remodeling of RDAPs

irCLIP-RNP was next used to study the dynamic remodeling of *cis*-RBP assemblies on RNA. The kinetic HNRNPC RDAP landscape was studied in response to EGF signaling. EGF was chosen as a stimulus for its well-known capacity to induce changes in RNA transcription, splicing, stability, and translation^22-24^ (**Fig. 3a**), while HNRNPC was selected as an RBP for its ability to bind and regulate numerous RNAs^25,26^. An EGF-stimulation timecourse was performed in EGF-responsive A431 cells followed by HNRNPC irCLIP-RNP (**Fig. 3b, Extended Data Fig. 6a-b**). Two RNP subzones ranging from 75-120kDa and 120-350kDa were subjected to mass spectrometry. Differential enrichment analysis identified 19 dynamic HNRNPC RDAPs across EGF stimulation (**Fig. 3c, Extended Data Fig. 6c, Supplementary Table 6**); these EGF-responsive RDAPs were mainly associated with mRNA splicing regulation (**Extended Data Fig. 6d-e**). Differences in the magnitude of the changes during EGF stimulation could also be observed between RNP subzones, indicating that the RDAP landscape of HNRNPC is dynamically remodeled in response to EGF stimuli (**Fig. 3c**). These data demonstrate that irCLIP-RNP captures dynamic RNA-RBP assemblies. As an additional illustration of this capacity, irCLIP-RNP was performed for the CAPRIN1 RBP after sodium arsenite-induced stress granule formation; this defined recruitment kinetics for a set of CAPRIN1 RDAPs distinct from those assembled with HNRNPC in response to EGF (**Extended Data Fig. 6f-j, Supplementary Table 7**). These data demonstrate diverse RDAPs can be dynamically recruited with specific RBPs in response to differing cell stimuli.

Protein-protein interaction analysis^27^ revealed an interaction network between HNRNPC and EGF-responsive RDAPs related to the regulation of mRNA processing, including canonical HNRNPC interactors^27^ (**Extended Data Fig. 6e**). Interestingly, a number of HNRNPC RDAPs showed reduced association on HNRNPC-bound RNAs upon EGF stimulation; in contrast, UPF1 and CSDE1 proteins joined HNRPNC on RNA in response to EGF (**Extended Data Fig. 6e**), suggesting that HNRNPC may rely on newly arrived co-bound RBPs to exert its EGF-responsive functions. To examine this hypothesis, RNA splicing was examined in EGF-treated A431 cells with and without depletion of HNRNPC, UPF1, or CSDE1 by RNA interference (**<u>RNAi</u>**) (**Extended Data Fig. 7a-c**). Splicing analysis was performed on the resulting RNA sequencing data to identify significant alternative splicing (**<u>AS</u>**) events (**Supplementary Table 8**). AS events were classified as alternative 3’ splice site (**<u>A3SS</u>**), alternative 5’ splice site (**<u>A5SS</u>**), exon skipping (**<u>SE</u>**), intron retention (**<u>RI</u>**), and mutually exclusive exons (**<u>MXE</u>**). SE events were the most common type of alternative splicing seen in all the tested RBPs, with depletion of HNRNPC itself displaying the greatest impacts (**Extended Data Fig. 7d**), consistent with its known biology^28,29^. Correlation analysis of significant AS events showed that, among those tested, HNRNPC displayed the highest similarity with UPF1 for RI transcripts and with CSDE1 for A3SS (**Fig. 3d**), suggesting that HNRNPC may act with discrete RDAPs to exert its pleiotropic impacts on RNA. Notably, the Δ percent of spliced-in (**<u>ΔPSI</u>**) value heatmaps and upset plots for each AS event category highlighted similarities between RI events altered by HNRNPC and UPF1 depletion (**Fig. 3e-f, Extended Data Fig. 7e**), suggesting joint action by HNRNPC and UPF1 in the regulation of intron-retained transcripts.

To further assess increased HNRNPC-UPF1 association after EGF signaling, proximity ligation assay (**<u>PLA</u>**) was performed using antibodies that recognize endogenous HNRNPC and UPF1 proteins. HNRNPC-UPF1 PLA signal was diminished after the knockdown of either HNRNPC or UPF1-KD, confirming its specificity (**Extended Data Fig. 7f-i**). Consistent with irCLIP-RNP data, HNRNPC-UPF1 proximity increased substantially during EGF stimulation in a time-dependent manner (**Fig. 3g-h**); this increased PLA signal was not due to elevated protein levels of HNRNPC or UPF1 (**Extended data Fig. 7j**). HNRNPC-UPF1 proximity signal increased proportionately more in the nucleus after 1h EGF treatment (**Fig. 3i**). The presence of PLA signal in both cytoplasm and nucleus suggests that HNRNPC and UPF1 actions might involve both cellular compartments. Indeed, subcellular fractionation showed that a small portion of UPF1 and HNRNPC proteins were also present in the nucleus or cytoplasm, respectively (**Extended data Fig. 7k-l**). Consistent with this possibility, and despite its largely nuclear localization, HNRNPC has been shown to translocate in the cytoplasm to influence cell proliferation^29,30^. Likewise, UPF1 has been shown to localize in both compartments, even interacting with chromatin^31,32^. These findings support irCLIP-RNP findings with HNRNPC and UPF1 and are consistent with the premise that EGF enhances the proximity of HNRNPC to UPF1 in multiple locations within the cell.

## UPF1 supports HNRNPC splicing surveillance of EGF-regulated proliferation mRNAs

UPF1 is an ATP-dependent RNA helicase component of the nonsense-mediated mRNA decay machinery (**<u>NMD</u>**), which recognizes and degrades aberrant mRNA transcripts^33^. The essential role of UPF1 in NMD raised the possibility that HNRNPC physically collaborates with UPF1 in surveilling the quality of EGF-stimulated transcripts to ensure the proper transduction of EGF signals. In support of this possibility, HNRNPC and UPF1 shared a significant portion of their RI events, namely 343, residing on 280 genes (**Extended Data Fig. 8a**). Notably, the 280 shared HNRNPC-UPF1 RI genes were enriched for those mediating cell cycle progression, response to external stimuli, and nucleic acid metabolism (**Extended Data Fig. 8b**), suggesting that HNRNPC and its UPF1 RDAP may collaborate in the regulation of mRNAs involved in biological processes induced by EGF signaling.

Next, we analyzed the expression and stability changes of the RNAs displaying the 343 shared HNRNPC and UPF1 RI events upon HNRNPC and UPF1 knockdown. We first performed differential transcript expression analysis of transcript abundances (**Supplementary Table 9**). This identified 186 RI transcripts to be significantly (q-value < 0.05) differentially expressed (**<u>DE</u>**) in both HNRNPC and UPF1 knockdown samples (**Extended Data Fig. 8c**). log_2_FC values of common DE RI transcripts showed significantly higher correlation than background differentially expressed transcripts between HNRNPC and UPF1, supporting the involvement of these two factors in splicing regulation (**Extended Data Fig. 8d**). Moreover, cumulative fraction analysis showed that common DE RI transcripts in HNRNPC or UPF1 knockdown data displayed significantly higher log_2_FC values than background differentially expressed transcripts (**Extended Data Fig. 8e**), suggesting that UPF1 and HNRNPC may collaborate in regulating the stability of EGF-responsive intron-retained transcripts. To test this, mRNA stability was assessed after the knockdown of HNRNPC and UPF1 in the context of EGF stimulation (**Extended Data Fig. 8f-h**). Principal component (**<u>PCA</u>**) analysis of mRNA stability showed distinct clustering of HNRNPC and UPF1 depleted cells for each time point in every sample after actinomycin D treatment (**Extended Data Fig. 8i**). Comparison of mRNA decay profiles between control and HNRNPC/UPF1 siRNA samples identified 158 RI transcripts that showed significant differential stability (**<u>DS</u>**) in both HNRNPC and UPF1 knockdown samples (**Supplementary Table 10**). Although the max(log_2_FC) values across actinomycin D treatment of the common DS RI transcripts did not display a higher correlation between HNRNPC and UPF1 knockdown samples than background DS transcripts, cumulative fraction analysis showed a significant tendency of common DS RI transcripts to have a higher log_2_FC (**Extended Data Fig. 8j-k**). Taken together, these data support a model where UPF1 and HNRNPC jointly control the expression and stability of intron-retained transcripts after EGF stimulation.

To explore whether HNRNPC and UPF1 influence each other’s binding on their intron-retained transcript targets, irCLIPv2 and Re-CLIP were performed at time points after EGF stimulation (**Extended Data Fig. 9a-c**), focusing on protein-coding genes because they comprised 95% of the 280 shared RI genes (**Extended Data Fig. 9d**). Most of the 307 RI regions jointly controlled by HNRNPC and UPF1 reside in UTRs (**Extended Data Fig. 9e**), where intron retention is known to influence mRNA stability^34^. Analysis of the distribution of the RT stops across UTRs, exon, and intron of the RI transcripts demonstrated that HNRNPC and UPF1 showed higher density in intronic and 3’UTR transcript regions, respectively (**Fig. 4a-b**), consistent with prior work^35,36^. Remarkably, Re-CLIP of HNRNPC followed by UPF1 showed a time-dependent significant higher density in the 3’UTR (**Fig. 4a-b**), suggesting that co-binding events between UPF1 and HNRNPC may occur in the 3’UTR of their co-regulated RNA targets.

To identify 3’UTR regions in RI transcripts responsive to EGF stimulation, correlation analysis was performed between the Z score values of total signal across the 3’UTR regions of RI transcripts of HNRNPC and UPF1 irCLIPv2 data. This demonstrated that HNRNPC and UPF1 increased their binding intensity at several 3’UTR regions, with the most significant correlation 60 minutes after EGF stimulation (**Fig. 4c**). Among these, 42 RI 3’UTR regions were selected with max(Z-score) > 0 across EGF stimulation. Overlap of the 35 co-bound genes of the 42 RI 3’UTR regions with the genes displaying either differential expression and/differential stability after HNRNPC and UPF1 knockdown defined 26 genes to be co-bound and co-regulated (**<u>co-BR</u>**) by HNRNPC and UPF1 in a time-dependent manner during EGF stimulation (**Fig. 4d and Extended Data Fig. 8l**). Minigene assay after HNRNPC and UPF1 knockdown of two co-BR genes, RND3 and RPS9, confirmed the splicing phenotype at these loci (**Extended Data Fig. 8m-o**). HNRNPC and UPF1 irCLIPv2 after knockdown of either UPF1 or HNRNPC at 0 and 60 minutes of EGF stimulation revealed that the binding of HNRNPC and UPF1 at the 30 EGF-responsive co-BR 3’UTR regions of the RI transcripts is highly dysregulated without a particular pattern (**Extended Data Fig. 9f-i**), suggesting a diverse set of interaction dependencies may occur between HNRNPC and UPF1 at these sites. A profile plot of the 30 EGF-responsive 3’UTR regions in Re-CLIP data showed that after 60 minutes of EGF, most of the signal was enriched near the 5’ ends of 3’UTRs (**Fig. 4e**). Normalized coverage heatmaps highlighted a subset of the 30 co-BR regions by HNRNPC and UPF1 in the 5’ of the 3’UTR with time-dependent increased intensity (**Fig. 4f**). Notably, de-novo motif analysis reported two poly(U)-containing motif that resembled the binding motif of HNRNPC (**Extended Data Fig. 9j**), further consistent with HNRNPC binding at these regions.

The top two 3’UTR regions displaying gained HNRNPC and UPF1 protein binding reside in the RND3 and DDX3X RNAs (**Fig. 4e-f**), both of which have known roles in EGF signaling^37,38^. RND3 and DDX3X RNAs were therefore chosen for further study. Bulk RNA sequencing of EGF-stimulated A431 cells demonstrated increased RND3 and DDX3X RNA expression, with RND3 upregulation comparable to the well-known early EGF-responsive genes FOSB and EGR1 (**Extended Data Fig. 9k-l, Supplementary Table 11**). Two prominent HNRNPC-UPF1 co-bound sites in the 3’UTRs of RND3 and DDX3X mRNAs were more closely examined with structural predictions using AlphaFold3^39^. Consistent with Re-CLIP data and de-novo motif analysis, structural modeling predicted HNRNPC and UPF1 proteins to interact with 3’UTR sequences in both the UUUUUUUUUUGUUA sequence of the RND3 RNA as well as with the 5’-UUG and UUUUG-3’ of the DDX3X RNA, respectively (**Fig 4g-h, Extended Data Fig. 9m-n**). Pull-down of in-vitro transcribed RNA previously incubated with EGF-treated A431 total cell lysate confirmed HNRNPC and UPF1 binding at RND3 and DDX3X 3’UTR regions (**Extended Data Fig. 9o-p**). These data support a model (**Fig. 4i**) in which HNRNPC and UPF1 influence each other’s binding at the 3’UTR regions of specific target RNAs to surveil the EGF-modulated RNA portfolio and indicate that irCLIP-RNP and Re-CLIP provide methods to study multiprotein assemblies on RNA molecules in living cells.

## Acknowledgments

We thank A.E. Oro, H.Y. Chang, and A. Fire for pre-submission review, M. Pilo, P. Bernstein, and A. Dazey for expert administrative assistance, and members of the Khavari lab for helpful discussions. This work was supported by NIAMS/NIH grants AR045192 and AR076965 to P.A.K. and by USVA Merit Review grant BX001409 to P.A.K., by R35 ES031707 to Y.W., by K01AR071481 to B.J.Z., by K99 through NIGMS K99GM147304 to N.M.R., and by a Swiss National Science Foundation Postdoc Mobility Fellowship P500BP-203019 to L.D.

## Author Contributions

B.J.Z., L.D., R.A.F., and P.A.K. conceived the project. L.D., B.J.Z., D.F.P., W.M., N.M.R. and L.V.J. performed experiments. L.D., B.J.Z., R.M.M., D.F.P., N.M.R, and S.S.. performed data analysis. Y.Y. and Y.W. supported with spectrometry data generation. R.A.F. and P.A.K. guided experiments and data analysis. L.D. and P.A.K. wrote the manuscript with input from all co-authors.

## References

1. Gebauer, F., Schwarzl, T., Valcárcel, J. & Hentze, M. W. RNA-binding proteins in human genetic disease. Nat. Rev. Genet. 22, 185–198 (2021).

2. Hentze, M. W., Castello, A., Schwarzl, T. & Preiss, T. A brave new world of RNA-binding proteins. Nat. Rev. Mol. Cell Biol. 19, 327–341 (2018).

3. Buccitelli, C. & Selbach, M. mRNAs, proteins and the emerging principles of gene expression control. Nat. Rev. Genet. 21, 630–644 (2020).

4. Mehta, M., Raguraman, R., Ramesh, R. & Munshi, A. RNA binding proteins (RBPs) and their role in DNA damage and radiation response in cancer. Adv. Drug Deliv. Rev. 191, 114569 (2022).

5. Heyn, L., Finset, A. & Ruland, C. M. Talking about feelings and worries in cancer consultations: the effects of an interactive tailored symptom assessment on source, explicitness, and timing of emotional cues and concerns. Cancer Nurs 36, E20–30 (2013).

6. Zarnegar, B. J., Flynn, R. A., Shen, Y., Do, B. T., Chang, H. Y. & Khavari, P. A. irCLIP platform for efficient characterization of protein-RNA interactions. Nat. Methods 13, 489–492 (2016).

7. Ramanathan, M., Porter, D. F. & Khavari, P. A. Methods to study RNA-protein interactions. Nat. Methods 16, 225–234 (2019).

8. Hafner, M., Katsantoni, M., Köster, T., Marks, J., Mukherjee, J., Staiger, D., Ule, J. & Zavolan, M. CLIP and complementary methods. Nature Reviews Methods Primers 1, 20 (2021).

9. Lee, F. C. Y. & Ule, J. Advances in CLIP Technologies for Studies of Protein-RNA Interactions. Mol. Cell 69, 354–369 (2018).

10. Schell, B., Legrand, P. & Fribourg, S. Crystal structure of SFPQ-NONO heterodimer. Biochimie 198, 1–7 (2022).

11. Wandrey, F., Montellese, C., Koos, K., Badertscher, L., Bammert, L., Cook, A. G., Zemp, I., Horvath, P. & Kutay, U. The NF45/NF90 Heterodimer Contributes to the Biogenesis of 60S Ribosomal Subunits and Influences Nucleolar Morphology. Mol. Cell. Biol. 35, 3491–3503 (2015).

12. Damianov, A., Ying, Y., Lin, C.-H., Lee, J.-A., Tran, D., Vashisht, A. A., Bahrami-Samani, E., Xing, Y., Martin, K. C., Wohlschlegel, J. A. & Black, D. L. Rbfox Proteins Regulate Splicing as Part of a Large Multiprotein Complex LASR. Cell 165, 606–619 (2016).

13. Choi, Y. D. & Dreyfuss, G. Isolation of the heterogeneous nuclear RNA-ribonucleoprotein complex (hnRNP): a unique supramolecular assembly. Proc. Natl. Acad. Sci. U.S.A. 81, 7471–7475 (1984).

14. Pandolfo, M., Valentini, O., Biamonti, G., Rossi, P. & Riva, S. Large-scale purification of hnRNP proteins from HeLa cells by affinity chromatography on ssDNA-cellulose. Eur J Biochem 162, 213–220 (1987).

15. Swanson, M. S. & Dreyfuss, G. Classification and purification of proteins of heterogeneous nuclear ribonucleoprotein particles by RNA-binding specificities. Mol. Cell. Biol. 8, 2237–2241 (1988).

16. Stark, C., Breitkreutz, B.-J., Reguly, T., Boucher, L., Breitkreutz, A. & Tyers, M. BioGRID: a general repository for interaction datasets. Nucleic Acids Res. 34, D535–9 (2006).

17. Geiger, T., Wehner, A., Schaab, C., Cox, J. & Mann, M. Comparative proteomic analysis of eleven common cell lines reveals ubiquitous but varying expression of most proteins. Mol Cell Proteomics 11, M111.014050 (2012).

18. Briata, P., Bordo, D., Puppo, M., Gorlero, F., Rossi, M., Perrone-Bizzozero, N. & Gherzi, R. Diverse roles of the nucleic acid-binding protein KHSRP in cell differentiation and disease. Wiley Interdiscip Rev RNA 7, 227–240 (2016).

19. Papadopoulou, C., Patrinou-Georgoula, M. & Guialis, A. Extensive association of HuR with hnRNP proteins within immunoselected hnRNP and mRNP complexes. Biochim. Biophys. Acta 1804, 692–703 (2010).

20. Porter, D. F., Miao, W., Yang, X., Goda, G. A., Ji, A. L., Donohue, L. K. H., Aleman, M. M., Dominguez, D. & Khavari, P. A. easyCLIP analysis of RNA-protein interactions incorporating absolute quantification. Nat Commun 12, 1569–16 (2021).

21. Feng, H., Bao, S., Rahman, M. A., Weyn-Vanhentenryck, S. M., Khan, A., Wong, J., Shah, A., Flynn, E. D., Krainer, A. R. & Zhang, C. Modeling RNA-Binding Protein Specificity In Vivo by Precisely Registering Protein-RNA Crosslink Sites. Mol. Cell 74, 1189–1204.e6 (2019).

22. Burgess, A. W. Regulation of Signaling from the Epidermal Growth Factor Family. J Phys Chem B 126, 7475–7485 (2022).

23. Nava, M., Dutta, P., Zemke, N. R., Farias-Eisner, R., Vadgama, J. V. & Wu, Y. Transcriptomic and ChIP-sequence interrogation of EGFR signaling in HER2+ breast cancer cells reveals a dynamic chromatin landscape and S100 genes as targets. BMC Med Genomics 12, 32–15 (2019).

24. Ma, H., Zhang, Z. & Tong, T. The effects of epidermal growth factor on gene expression in human fibroblasts. In Vitro Cell Dev Biol Anim 38, 481–486 (2002).

25. Geuens, T., Bouhy, D. & Timmerman, V. The hnRNP family: insights into their role in health and disease. Hum Genet 135, 851–867 (2016).

26. Mo, L., Meng, L., Huang, Z., Yi, L., Yang, N. & Li, G. An analysis of the role of HnRNP C dysregulation in cancers. Biomark Res 10, 19–10 (2022).

27. Szklarczyk, D., Gable, A. L., Lyon, D., Junge, A., Wyder, S., Huerta-Cepas, J., Simonovic, M., Doncheva, N. T., Morris, J. H., Bork, P., Jensen, L. J. & Mering, von, C. STRING v11: Protein-protein association networks with increased coverage, supporting functional discovery in genome-wide experimental datasets. Nucleic Acids Res. 47, D607–D613 (2019).

28. Zhou, R., Park, J. W., Chun, R. F., Lisse, T. S., Garcia, A. J., Zavala, K., Sea, J. L., Lu, Z.-X., Xu, J., Adams, J. S., Xing, Y. & Hewison, M. Concerted effects of heterogeneous nuclear ribonucleoprotein C1/C2 to control vitamin D-directed gene transcription and RNA splicing in human bone cells. Nucleic Acids Res. 45, 606–618 (2017).

29. Martino, F., Varadarajan, N. M., Perestrelo, A. R., Hejret, V., Durikova, H., Vukic, D., Horvath, V., Cavalieri, F., Caruso, F., Albihlal, W. S., Gerber, A. P., O’Connell, M. A., Vanacova, S., Pagliari, S. & Forte, G. The mechanical regulation of RNA binding protein hnRNPC in the failing heart. Sci Transl Med 14, eabo5715 (2022).

30. Kim, J. H., Paek, K. Y., Choi, K., Kim, T.-D., Hahm, B.Kim, K.-T. & Jang, S. K. Heterogeneous nuclear ribonucleoprotein C modulates translation of c-myc mRNA in a cell cycle phase-dependent manner. Mol. Cell. Biol. 23, 708–720 (2003).

31. Hong, D., Park, T. & Jeong, S. Nuclear UPF1 Is Associated with Chromatin for Transcription-Coupled RNA Surveillance. Mol. Cells 42, 523–529 (2019).

32. Singh, A. K., Choudhury, S. R., De, S., Zhang, J., Kissane, S., Dwivedi, V., Ramanathan, P., Petric, M., Orsini, L., Hebenstreit, D. & Brogna, S. The RNA helicase UPF1 associates with mRNAs co-transcriptionally and is required for the release of mRNAs from gene loci. Elife 8, (2019).

33. Kim, Y. K. & Maquat, L. E. UPFront and center in RNA decay: UPF1 in nonsense-mediated mRNA decay and beyond. RNA 25, 407–422 (2019).

34. Jacob, A. G. & Smith, C. W. J. Intron retention as a component of regulated gene expression programs. Hum Genet 136, 1043–1057 (2017).

35. Hurt, J. A., Robertson, A. D. & Burge, C. B. Global analyses of UPF1 binding and function reveal expanded scope of nonsense-mediated mRNA decay. Genome Research 23, 1636–1650 (2013).

36. Xing, S., Wang, J., Wu, R., Hefti, M. M., Crary, J. F. & Lu, Y. Identification of HnRNPC as a novel Tau exon 10 splicing factor using RNA antisense purification mass spectrometry. RNA Biol 19, 104–116 (2022).

37. Almarán, B., Ramis, G., Fernández de Mattos, S. & Villalonga, P. Rnd3 Is a Crucial Mediator of the Invasive Phenotype of Glioblastoma Cells Downstream of Receptor Tyrosine Kinase Signalling. Cells 11, 3716 (2022).

38. Nozaki, K., Kagamu, H., Shoji, S., Igarashi, N., Ohtsubo, A., Okajima, M., Miura, S., Watanabe, S., Yoshizawa, H. & Narita, I. DDX3X induces primary EGFR-TKI resistance based on intratumor heterogeneity in lung cancer cells harboring EGFR-activating mutations. PLoS ONE 9, e111019 (2014).

39. Abramson, J., Adler, J., Dunger, J., Evans, R., Green, T., Pritzel, A., Ronneberger, O., Willmore, L., Ballard, A. J., Bambrick, J., Bodenstein, S. W., Evans, D. A., Hung, C.-C., O’Neill, M., Reiman, D., Tunyasuvunakool, K., Wu, Z., Žemgulytė, A., Arvaniti, E., Beattie, C., Bertolli, O., Bridgland, A., Cherepanov, A., Congreve, M., Cowen-Rivers, A. I., Cowie, A., Figurnov, M., Fuchs, F. B., Gladman, H., Jain, R., Khan, Y. A., Low, C. M. R., Perlin, K., Potapenko, A., Savy, P., Singh, S., Stecula, A., Thillaisundaram, A., Tong, C., Yakneen, S., Zhong, E. D., Zielinski, M., Žídek, A., Bapst, V., Kohli, P., Jaderberg, M., Hassabis, D. & Jumper, J. M. Accurate structure prediction of biomolecular interactions with AlphaFold 3. Nature 630, 493–500 (2024).

