## Supplementary Information for "irCLIP-RNP and Re-CLIP reveal patterns of dynamic protein associations on RNA"

### Supplementary Discussion

irCLIP-RNP and Re-CLIP provide approaches to analyzing relationships between an RBP of interest and its *cis*-RDAP portfolio. irCLIP-RNP on a panel of RBPs in two cell types showed the existence of selective patterns of RBP-specific *cis*-binding RDAP relationships in differing cell type contexts, further supported by co-binding analysis with Re-CLIP. Re-CLIP defined regions that were simultaneously co-bound by both the RBP tested and the identified RDAPs. An irCLIP-RNP and Re-CLIP time course after EGF stimulation quantified RBP-RDAP remodeling kinetics on co-bound RNAs, uncovering EGF-induced recruitment of UPF1 on HNRNPC-bound RNA to mediate RNA splicing surveillance, notably at 3'UTRs. Taken together, these data point to the possible existence of a patterned set of cell-context modulated combinatorial RBP "codes" co-bound to RNA that may be analogous to cell-type selective combinations of transcription factors that bind regulatory elements in DNA.

Compared to a prior CLIP-based method coupled to MS[1], irCLIP-RNP uses specific on-bead RNA ligation of the infrared biotinylated adapter and streptavidin pull-down to specifically enrich proteins bound in *cis* to the same RNAs that are bound by the immunoprecipitated RBP of interest. This unique irCLIP-RNP feature marks ligated RNPs with an infrared dye to monitor, in real-time, the profiles of different RNPs across a denaturing gel. irCLIP-RNP enables the classification of RDAPs according to their migratory pattern on denaturing gels, providing clues about the degree of multimerization of those associations in different cellular conditions. For example, a cell-type selective connection between HNRNPU, NONO, and SFPQ was observed in which these proteins co-immunoprecipitate on RNA at different MW bands depending on the cell type. Such findings indicate that these factors assemble on RNAs influencing their RDAP interactome reciprocally, providing a rationale for further study of their functional impacts in different cellular contexts. An additional advantage of irCLIP-RNP is its facilitation of direct interrogation of these larger MW RNP complexes. By comparing the RDAP landscape of hnRNP proteins after digestion with different RNase doses, irCLIP-RNP uncovered RNA-dependent protein associations occurring at different RNA distances depending on the immunoprecipitated RBP.

The study of RNA-protein assemblies after EGF treatment highlights the capacity of irCLIP-RNP and Re-CLIP to characterize dynamic RDAP landscapes. The connection observed here between EGF treatment and HNRNPC regulation of EGF-induced mRNAs is also of interest, given the roles of EGF signaling in multiple human cancers[2]. HNRNPC has been observed to be upregulated in human cancers, where its overexpression has been linked to poor prognosis[3-6]. A role for HNRNPC in mediating EGF target gene expression, however, has not, to our knowledge, been previously described, providing impetus for additional similar studies of RNA-protein assemblies in response to an array of oncogenic cues. irCLIP-RNP uncovered an RNA-dependent association between HNRNPC and the NMD core factor, UPF1, in ensuring the proper splicing output after activation of EGF signaling. This is congruent with previous data where HNRNPC and UPF1 have been shown to regulate splicing events of Alu-element-containing exons in the nucleus (HNRNPC) or the cytoplasm (UPF1)[7]. Here, through integration of differential transcript and stability analysis with irCLIPv2 and Re-CLIP data, we provide evidence suggesting that activation of EGF signaling led to the physical co-binding of UPF1 to 3'UTR sites of HNRNPC RNA binding in intron-retained transcripts, controlling their fate. Specifically, HNRNPC and UPF1 proved essential in degrading incompletely spliced mRNAs induced by EGF stimulation, including mRNAs encoding proteins, such as RND3, involved in the regulation of cell proliferation, migration, invasion and stress response, which are essential outputs of EGF signaling[8]. For instance, UPF1 and HNRNPC depletion modulated each other's binding at 3'UTR regions within the regulated RND3 mRNA target, a downstream effector of EGF signaling through cytoskeleton regulation, where they were both required for the degradation of intron-retained transcripts induced after EGF treatment. irCLIP-RNP thus uncovered a role for HNRNPC and UPF1 in ensuring proper fidelity of mRNAs induced by EGF, thereby linking HNRNPC to a mediator of NMD.

### Extended Data Figure 1: irCLIP-RNP reveals distinct protein associations after intermediate RNase digestion

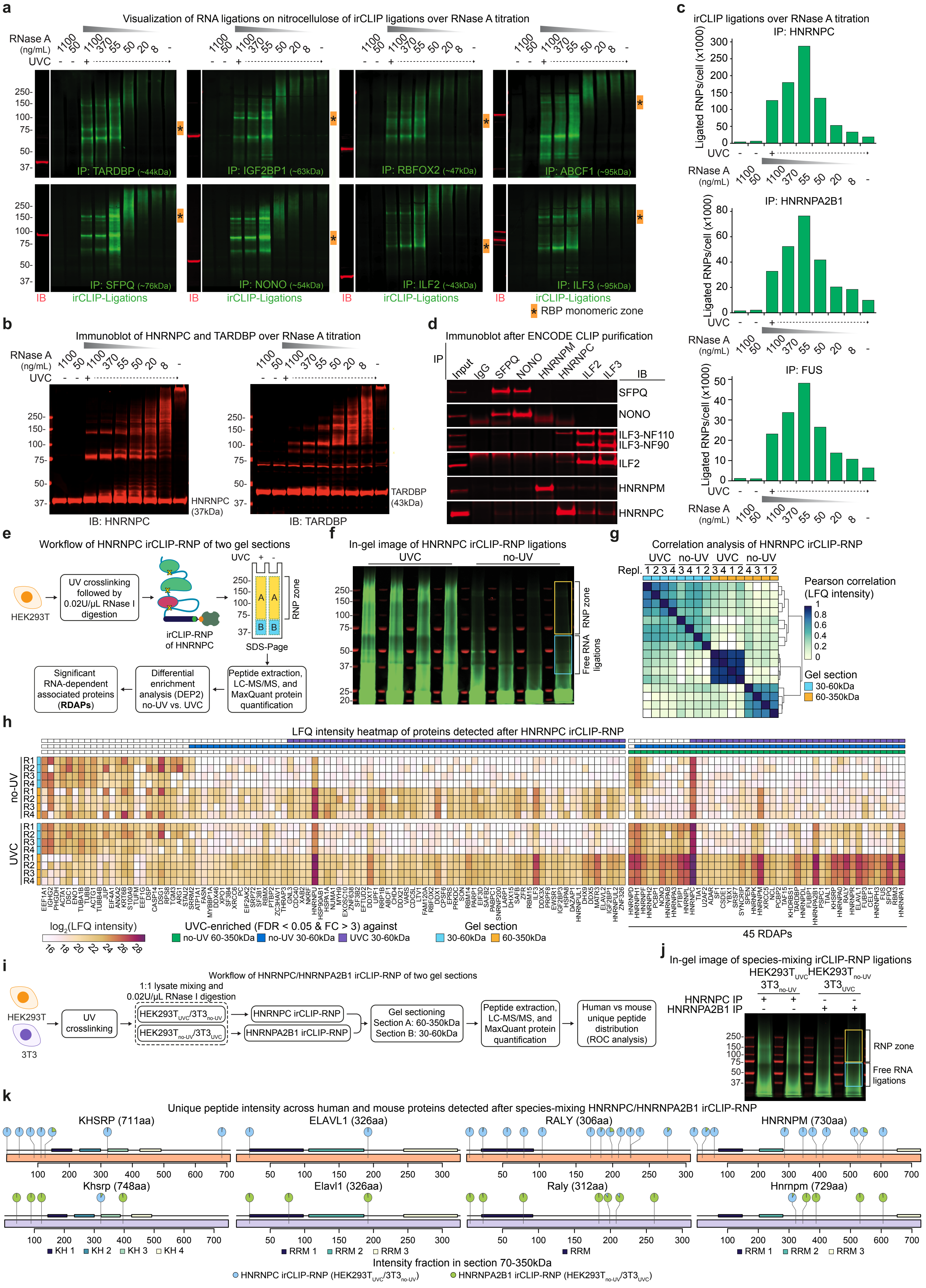

**Extended Data Figure 2: irCLIP-RNP identifies proximal and distal cis-RNA-dependent protein associations (RDAPs)**

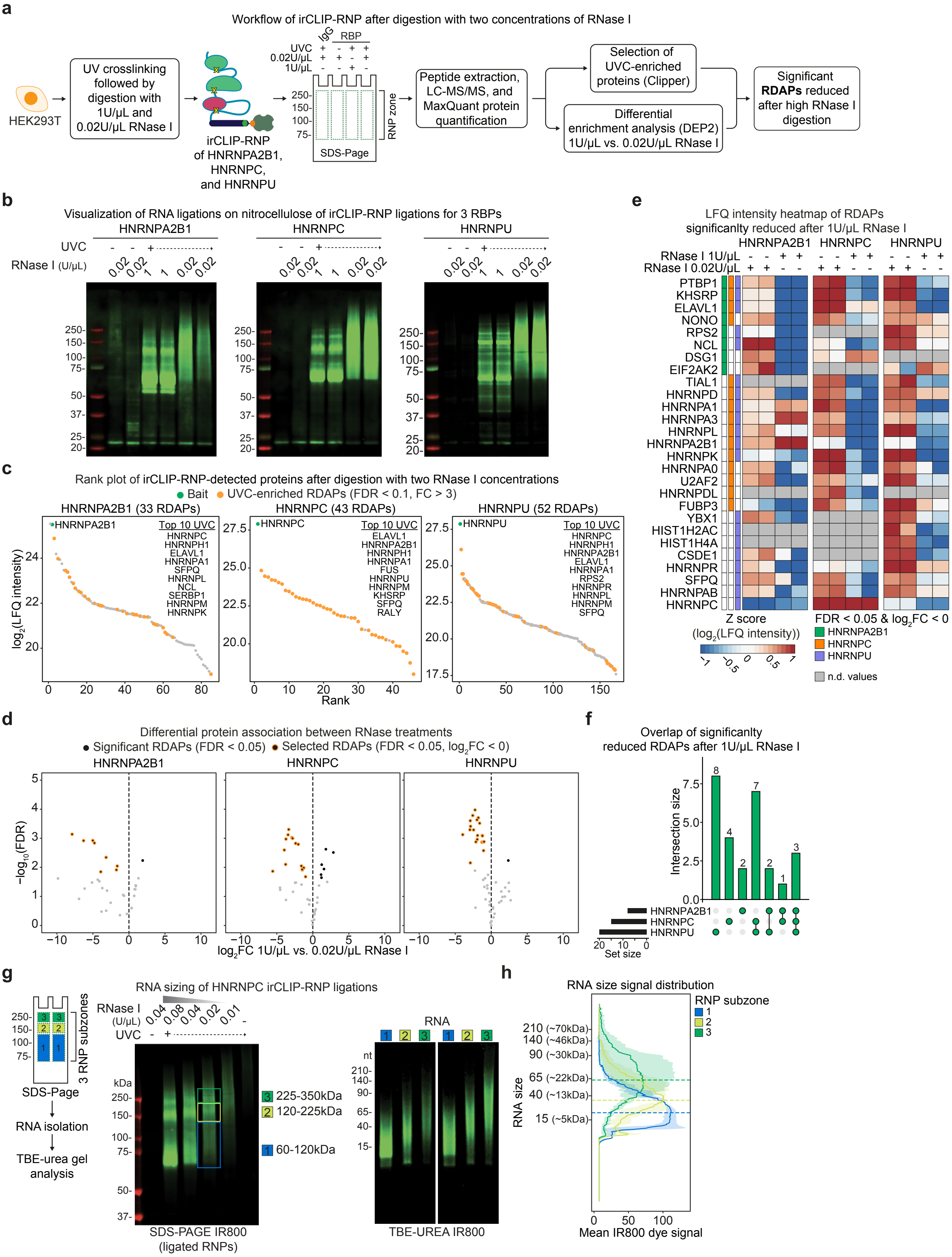

**Extended Data Figure 3: RBPs show different RDAP landscape between HEK293T and HepG2**

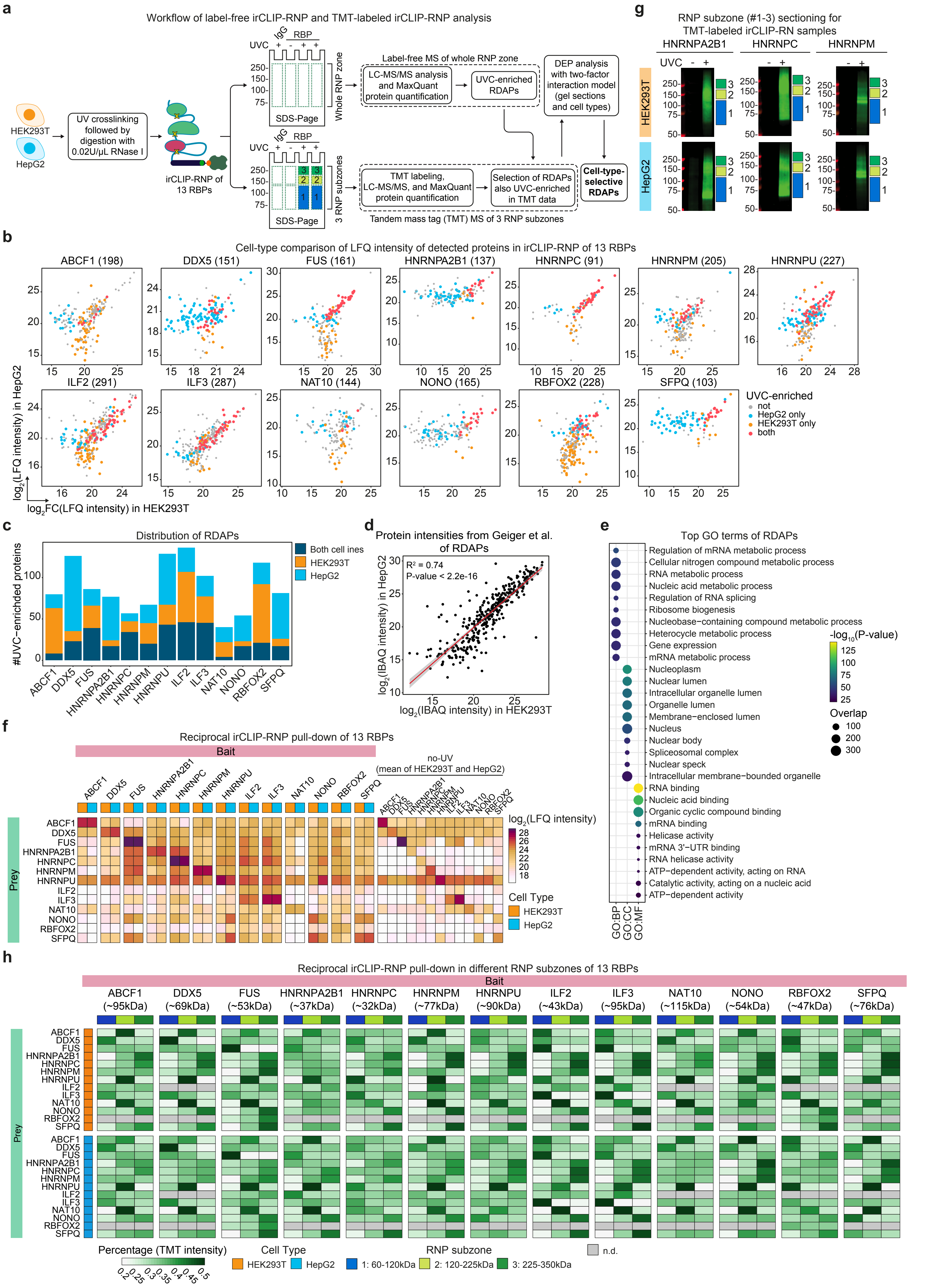

### Extended Data Figure 4: Cell-context-dependent RDAP assemblies of 13 RBPs

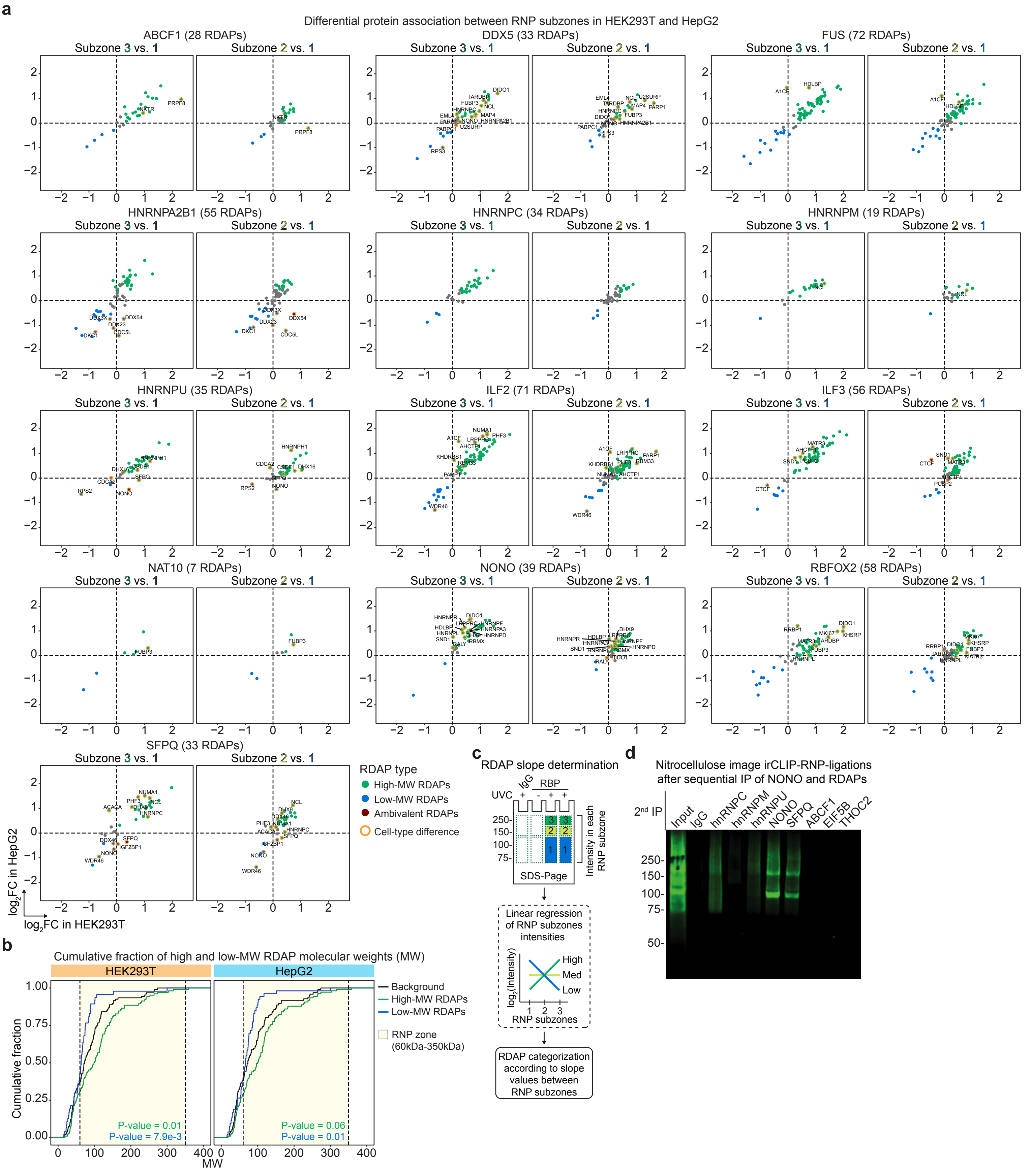

**Extended Data Figure 5: irCLIPv2 and Re-CLIP supports multi-RBP model on the same RNA molecule**

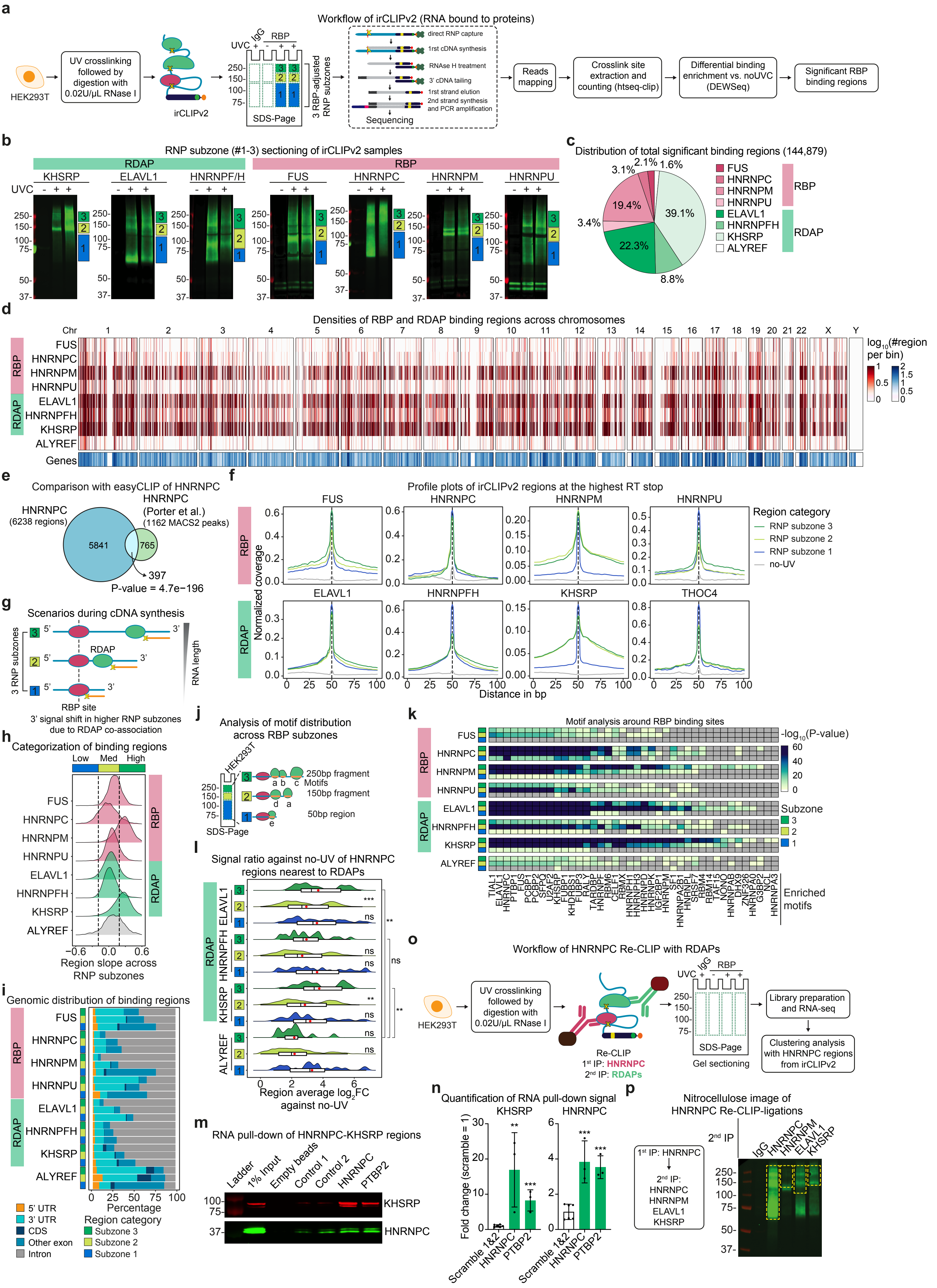

**Extended Data Figure 6: Dynamic remodeling of RDAP landscapes after cellular stimulations**

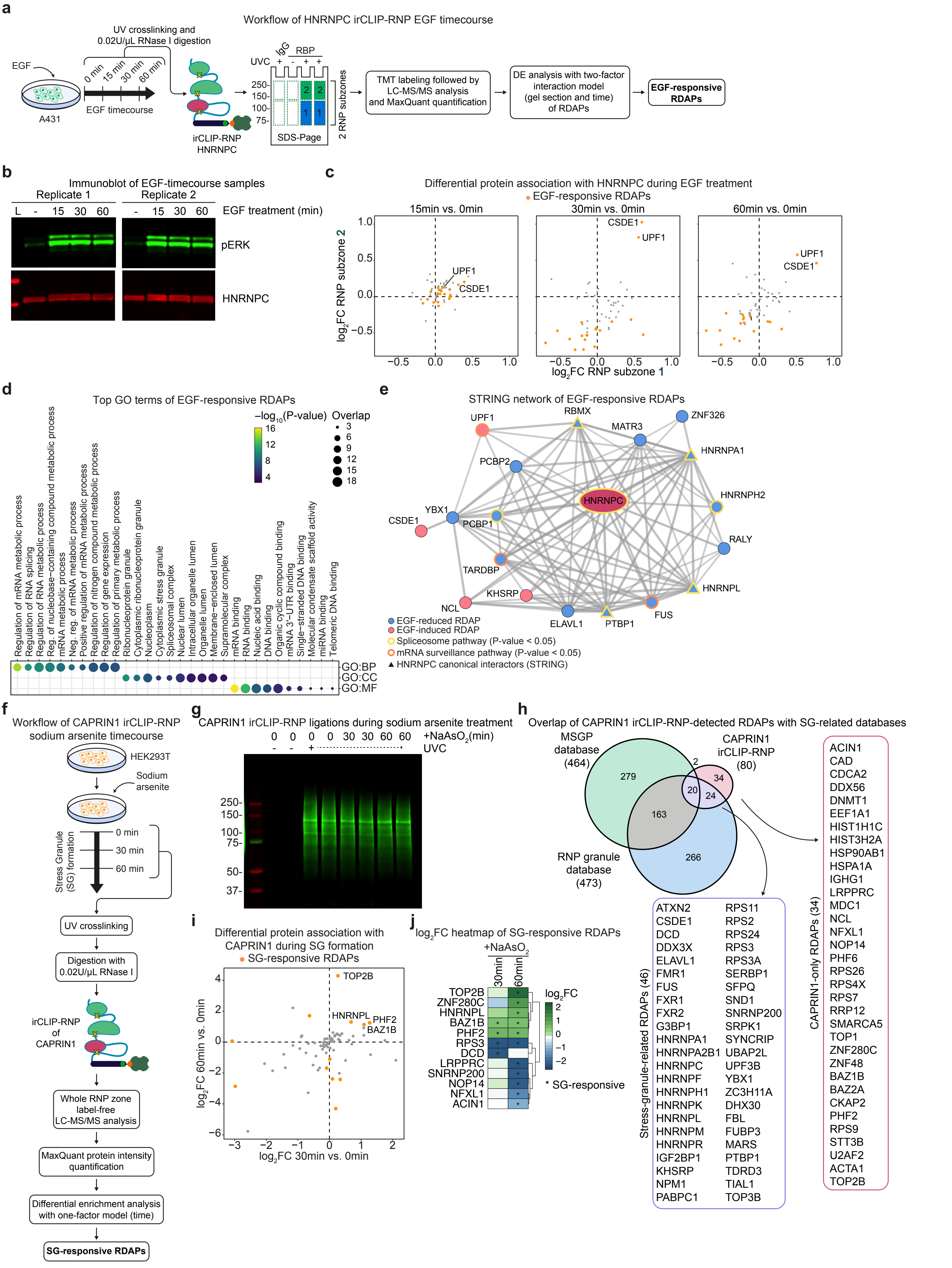

### Extended Data Figure 7: Splicing analysis of HNRNPC and RDAP-depleted A431 cells

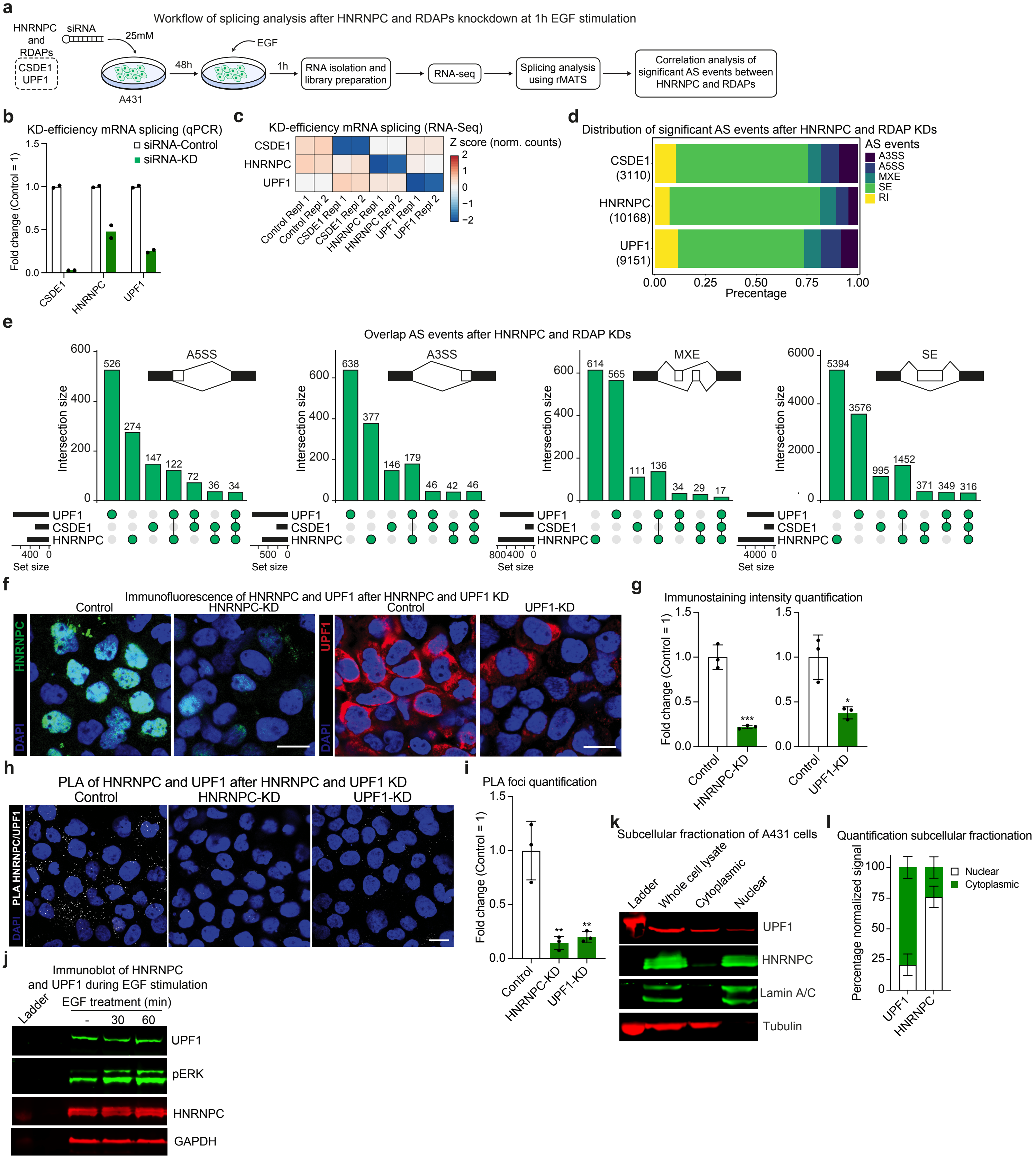

### Extended Data Figure 8: UPF1 and HNRNPC regulates the stability of RI RNA transcripts during EGF stimulation

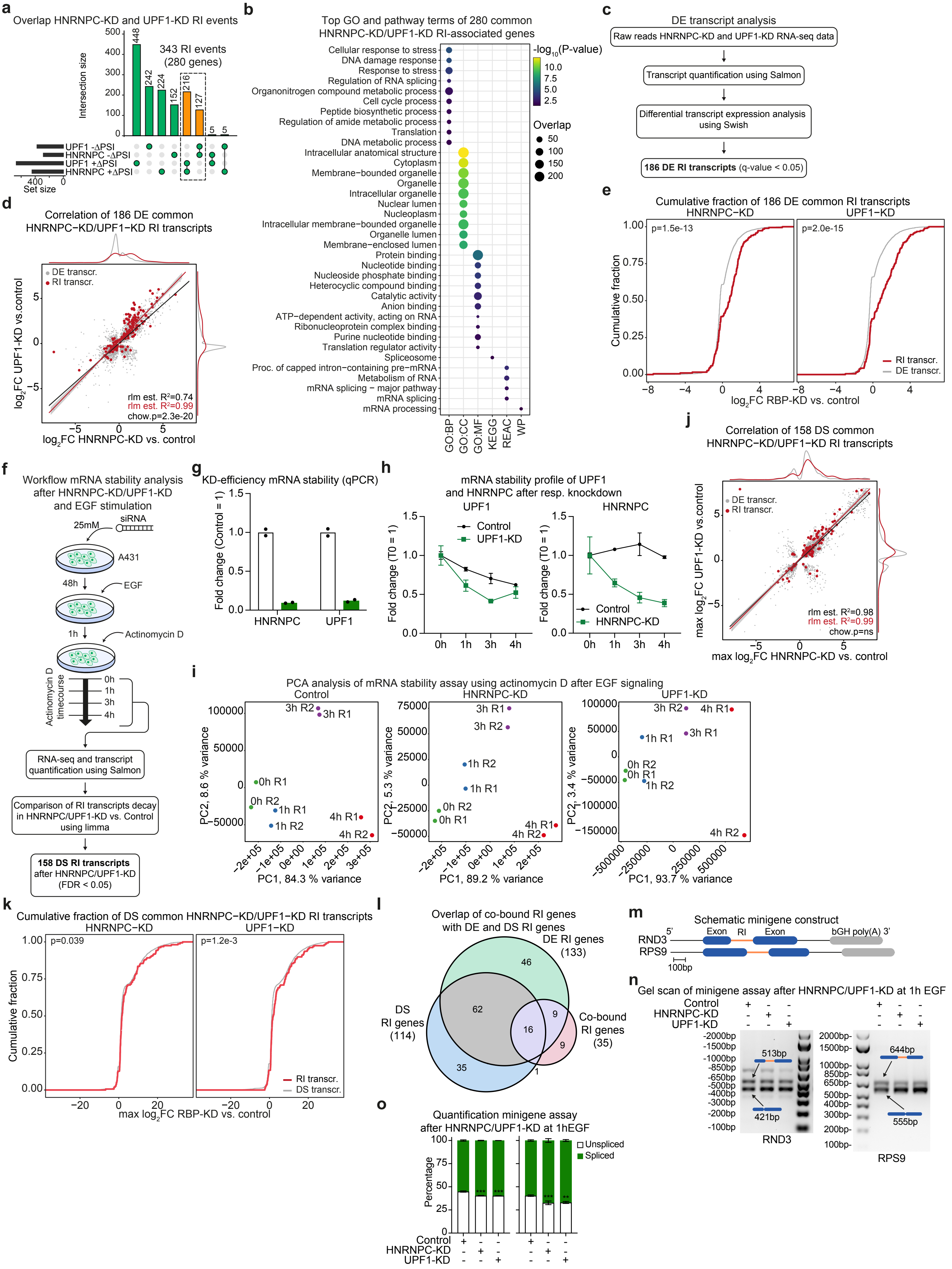

**Extended Data Figure 9: UPF1 supports HNRNPC RNA splicing surveillance of EGF-regulated mRNAs**

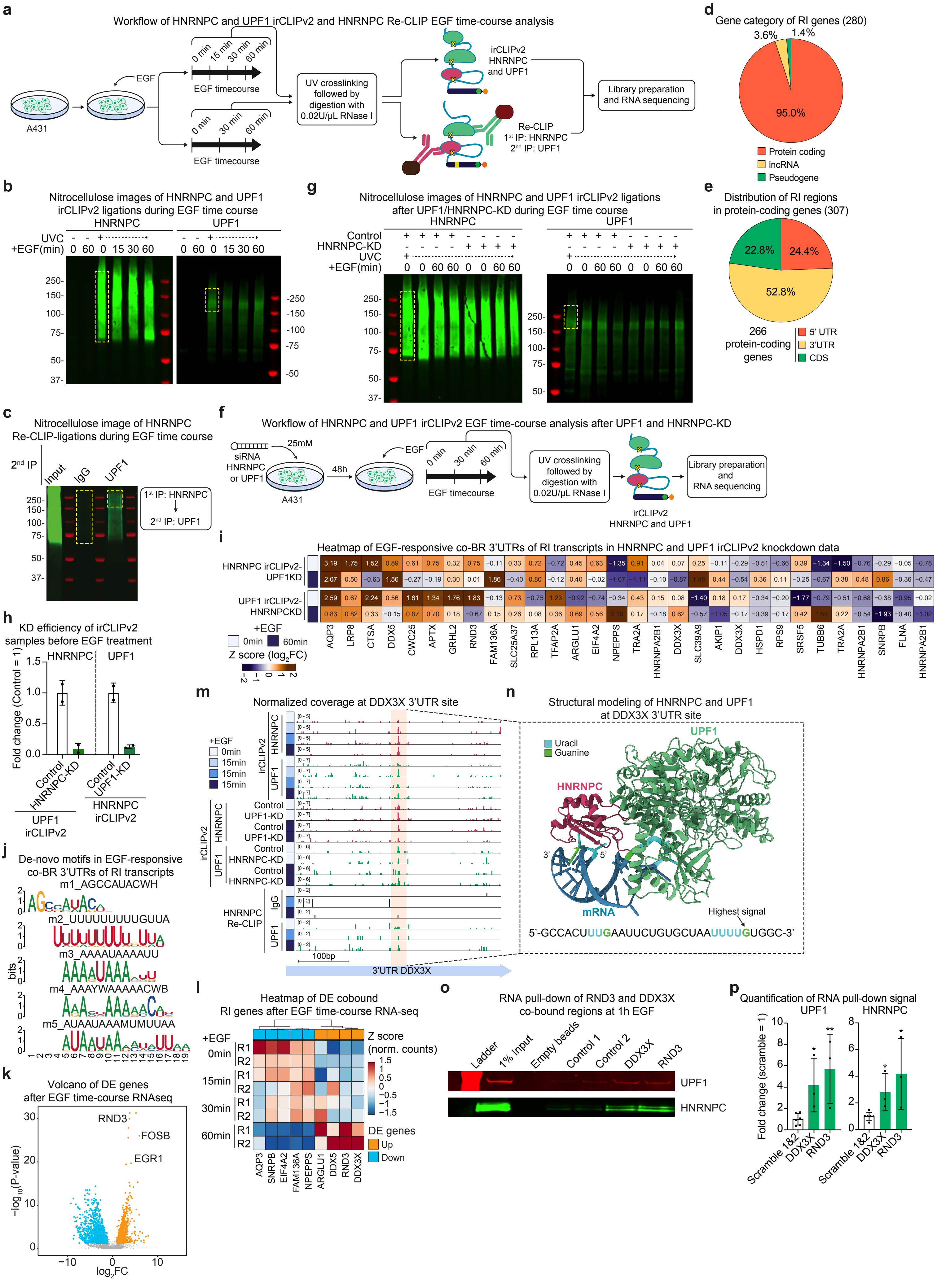

### Extended Data Figure Legends

**Extended Data Figure 1: irCLIP-RNP reveals distinct protein associations after intermediate RNase digestion. (a)** Nitrocellulose images indicate the infrared signal of irCLIP ligations subsequent to in-lysate digestion with six RNase A concentrations (from 1100 ng/mL to 8 ng/mL) for additional RBPs also involved in RNA processing. Green signal: irCLIP ligation; red signal: immunoblotting for the tested RBP; asterisks in orange box: RBP monomeric zone. The reported kDa for each RBP represents the calculated MW. **(b)** Immunoblot images indicate HNRNPC and TARDBP protein signal after in-lysate RNase A titration (1100ng/mL - 8ng/mL) followed by irCLIP ligation. **(c)** Bars indicate irCLIP ligations per cell calculated from infrared signal after dot plotting irCLIP ligations resulting from $1x10^4$ cells for HNRNPC, HNRNPA2B1, and FUS over six RNase A concentrations ranging from 1100 ng/mL to 8 ng/mL. UVC: UV cross-linking. **(d)** Immunoblot images indicate protein signal for SFPQ, NONO, ILF2/3, HNRNPM, and HNRNPC after immunoprecipitation following standard ENCODE CLIP purification conditions for SFPQ, NONO, HNRNPM, HNRNPC, ILF2, and ILF3 in HEK293T cells. IP: immunoprecipitated protein; IB: immunoblotted protein. **(e)** Workflow of HNRNPC irCLIP-RNP of two gel sections ranging from 30-60kDa and 60-350kDa in HEK293T cells (n = 4). **(f)** In-gel image indicates the infrared signal of HNRNPC irCLIP-RNP ligations for 4 UVC and 4 no-UV samples. Blue box: gel section corresponding to free RNA ligations; yellow box: gel section corresponding to the RNP zone. **(g)** Heatmap indicates Pearson correlation coefficients between 4 no-UV and 4 UVC HNRNPC irCLIP-RNP samples. Blue box: free RNA ligations gel section; yellow box: RNP zone gel section. **(h)** Heatmap indicates imputed $log_2$-transformed LFQ intensities of proteins detected after HNRNPC irCLIP-RNP. Green, blue, and purple boxes: significant proteins against no-UV 60-350kDa, no-UV 30-60kDa, and UVC 30-60kDa gel sections. **(i)** Workflow of HNRNPC/HNRNPA2B1 irCLIP-RNP species-mixing experiment using a 1:1 mixing of human (HEK293T) and mouse (3T3) cells (n = 2). **(j)** In-gel scan indicates the infrared signal of species-mixing irCLIP-RNP samples. **(k)** Pie charts indicate the ratio of imputed LFQ intensities of cell-type-specific unique peptides between HNRNPC irCLIP-RNP (human UVC; mouse no-UV) and HNRNPA2B1 irCLIP-RNP (human no-UV; mouse UVC) across KHSRP/Khsrp, ELAVL1/Elavl1, RALY/Raly, and HNRNPM/Hnrnpm proteins.

**Extended Data Figure 2: irCLIP-RNP identifies proximal and distal cis-RNA-dependent protein associations (RDAPs). (a)** Workflow of irCLIP-RNP to analyze the effects on RNA-dependent protein associations of 1U/µL and 0.02U/µL RNase I treatments in HEK293T (n = 2). **(b)** Nitrocellulose images indicate the infrared signal of irCLIP-RNP ligations after 1U/µL and 0.02U/µL RNase I digestions in HEK293T for HNRNPC, HNRNPA2B1, and HNRNPU. **(c)** Rank plots indicate imputed log2-transformed LFQ intensities of proteins detected after HNRNPA2B1/HNRNPC/HNRNPU irCLIP-RNP using two doses of RNase I (1U/µL and 0.02U/µL). Orange dot: UVC-enriched proteins in at least one RNase I dose. **(d)** Volcano plots indicate -$\log_{10}$(FDR) and $\log_2$FC values of differential RDAPs between 1U/µL and 0.02U/µL RNase I samples associated with HNRNPC, HNRNPA2B1, and HNRNPU. Black dots: significant proteins; orange circle: selected RDAPs with a significant reduction in 1U/µL RNase I treatment. **(e)** Heatmap indicate Z scores of imputed $\log_2$-transformed LFQ intensities of 1U/µL and 0.02U/µL RNase I samples for the 27 significant RDAPs associating with HNRNPC, HNRNPA2B1, and HNRNPU. Green, orange, and purple boxes: significant RDAPs for HNRNPA2B1, HNRNPC, and HNRNPU, respectively; grey boxes: n.d. values. **(f)** Bars indicate the overlap between HNRNPC, HNRNPA2B1, and HNRNPU of the 27 significantly reduced RDAPs after 1U/µL RNase I in-lysate digestion. **(g)** RNA sizing through TBE-Urea gel of HNRNPC irCLIP-RNP isolated RNA in HEK293T from the three RNP subzones ranging from 60-120kDa (blue), 120-225kDa (light green), and 225-350kDa (green). **(h)** Lines indicate mean infrared signal distribution of RNA isolated from HNRNPC irCLIP-RNP ligations from three RNP subzones shown in **g** (n = 2). Data are mean ± s.d. Blue line: 60-120kDa; light-green line: 120-225kDa; green line: 225-350kDa.

**Extended Data Figure 3: RBPs show different RDAP landscape between HEK293T and HepG2. (a)** Workflow of the integrated analysis of label-free MS from the whole RNP zone and the TMT analysis from the three RNP subzones for 13 RBPs in HEK293T and HepG2 (n = 2). **(b)** Scatter plots indicate imputed $\log_2$-transformed LFQ intensities in HEK293T and HepG2 of irCLIP-RNP-detected proteins across the 13 RBPs. Orange and blue dots: UVC-enriched only in HEK293T or HepG2, respectively; red dots: UVC-enriched in HEK293T and HepG2. **(c)** Stacked bars indicate the distribution of 360 RDAPs across the 13 RBPs in HEK293T and HepG2. **(d)** Scatter plot indicate imputed $\log_2$-transformed iBAQ intensities taken from Geiger et al.[9] in HEK293T and HepG2 of 346 out of the 360 RDAPs. Red line is fitted to the points using linear regression. **(e)** Bubble plot indicates -$log_{10}$(P-values) of top GO terms for cellular components, biological processes, and molecular functions of the 360 RDAPs. **(f)** Reciprocal heatmap indicates imputed $log_2$-transformed LFQ intensities between the 13 RBPs (dark fuchsia: bait; green: prey). LFQ intensities from no-UV samples are displayed as the mean of HEK293T and HepG2 samples. Orange boxes: HEK293T; light blue boxes: HepG2. **(g)** Nitrocellulose images indicate the infrared signal of TMT irCLIP-RNP ligations for the HNRNPA2B1, HNRNPC, and HNRNPM in HEK293T (orange) and HepG2 (blue). Blue, light-green, and green boxes highlight the gel sectioning of the three RNP subzones. **(h)** Reciprocal heatmap indicates the TMT intensity percentages across the three RNP subzones between the 13 RBPs (dark fuchsia: bait; green: prey) in HEK293T (orange) and HepG2 (blue). Blue, light-green, and green boxes: RNP subzones #1-3.

**Extended Data Figure 4: Cell-context-dependent RDAP assemblies of 13 RBPs.** **(a)** Volcano plots indicate -$log_{10}$(FDR) and $log_2$FC values of differentially enriched RDAPs with the 13 RBPs after comparing RNP subzones #2 and #3 against #1 (left and right plots, respectively). Green and blue dots: low and high-MW RDAPs; dark red dots: ambivalent distributed RDAPs between HEK293T and HepG2; orange circle: RDAPs with significant differences in TMT intensity distribution across the RNP subzones between HEK293T and HepG2. **(b)** Lines indicate the cumulative fraction of MWs of low and high-MW RDAPs and background proteins in HEK293T (left) and HepG2 (right). Black line: background proteins; green and blue lines: high and low-order RDAPs in HEK293T and HepG2. P-values between high and low-order RDAPs against the background in each cell line were calculated by performing the Kolmogorov-Smirnov test. **(c)** Schematic of the RDAP categorization in the different RNP subzones, according to slope values calculated between the RNP subzone #1-3 intensity ratios. **(d)** Nitrocellulose image indicates the infrared signal of sequential irCLIP-RNP pull-downs of NONO followed by the RDAPs HNRNPC, HNRNPM, HNRNPU, NONO, SFPQ, and ABCF1 as well as the negative control proteins EIF5B and THOC2.

**Extended Data Figure 5: irCLIPv2 and Re-CLIP support multi-RBP model on the same RNA molecule. (a)** Workflow of irCLIPv2 experiments of three RNP subzones for 3 RDAPs and 4 RBPs belonging to HNRNPs protein family in HEK293T (n = 2). **(b)** Nitrocellulose images indicate the infrared signal of irCLIP-RNP ligations for the 3 RDAPs (green) and 4 RBPs (dark fuchsia) in HEK293T. Blue, light-green, and green boxes highlight the gel sectioning of the three RNP subzones. **(c)** Pie chart indicates the percentages of total significant binding regions of 3 RDAPs (green) and 4 RBPs (dark fuchsia). **(d)** Heatmap indicates the number of significant regions per bin along each chromosome for RDAPs (green) and RBPs (dark fuchsia). Genes distribution across chromosomes are displayed in shades of blue. **(e)** Venn diagram indicates the overlap between irCLIPv2-identified regions for HNRNPC and previously published binding regions identified with easyCLIP[10]. **(f)** Lines indicate the normalized coverage at significant binding regions in the three RNP subzones and no-UV samples (blue: #1; light-green: #2; green: #3; grey: no-UV) for RBPs (dark fuchsia) and RDAPs (green). Coverage on the negative strand was reversed to be represented in a 5' to 3' orientation. An extension of +/- 50bp was applied from the highest RBP peaks in the RNP subzone #1. **(g)** Schematic representation of cDNA synthesis early truncation resulting in signal 3'-shift due to RDAPs co-binding. **(h)** Lines indicate the slope density distribution calculated from average normalized RT stop counts of significant regions for RDAPs (green) and RBPs (dark fuchsia). One standard deviation (s.d.) was used to classify the significant regions according to their occurrence in the three RNP subzones. Blue/light-green/green boxes: RNP subzones #1-3. **(i)** Stacked bars indicated the percentage of significant regions in transcriptomic landscape features (5' UTR, 3' UTR, CDS, other exons, and intron) for RBPs (green) and RDAPs (dark fuchsia). Blue/light-green/green boxes: RNP subzones #1-3. **(j)** Schematic representation of RDAPs motif analysis across RNP subzones in the regions directly downstream of highest RBP peaks. 250bp, 150bp, and 75bp fragments were used as input in the AME tool for RNP subzones #1 to #3, respectively. **(k)** Heatmap indicates the enriched motif P-values of high-MW proteins in the three RNP subzones of RDAPs (green) and RBPs (dark fuchsia). Blue/light-green/green boxes: RNP subzones #1-3. **(l)** Raincloud plots indicate the average $\log_2$FC against no-UV samples for HNRNPC regions categorized as low (blue), medium (light green), or high (green) distances in their corresponding RNP subzone. **P-value < 0.01, ***P-value < 0.001, ns not significant – ordinary one-way ANOVA with Tukey's test. Only the significance of high-categorized regions against low and medium categories related to the same protein or against the ALYREF long-distance category is shown. **(m)** Immunoblot image indicates the protein signal of HNRNPC and KHSRP after native RNA pull-down of biotinylated HNRNPC-KHSRP flanking regions in HEK293T cells. **(n)** Bars indicate the fold change against scramble control of HNRNPC and KHSRP protein signal after native RNA pull-down of biotinylated HNRNPC-KHSRP flanking regions in HEK293T cells. **P-value < 0.01, ***P-value < 0.001 – unpaired one-tailed Student's t-test against scramble control (n = 3-6). Data are mean ± s.d. **(o)** Workflow of Re-CLIP experiments where sequential immunoprecipitations of HNRNPC followed by the RDAPs ELAVL1, KHSRP, and HNRNPM were performed (n = 2). **(p)** Nitrocellulose image indicates the infrared signal of HNRNPC Re-CLIP ligations after sequential immunoprecipitation of itself and the RDAPs HNRNPM, ELAVL1, KHSRP, as well as the negative control IgG.

**Extended Data Figure 6: Dynamic remodeling of the RDAP landscape after cellular stimulations.** **(a)** Workflow of HNRNPC irCLIP-RNP-TMT during EGF stimulation in A431 cells (n = 2). **(b)** Immunoblot images indicate the phosphorylated ERK and HNRNPC protein signal in EGF-stimulated irCLIP-RNP-TMT samples. **(c)** Scatterplot indicates the RNP subzone #1-2 $\log_2$FC values of differentially enriched RDAPs during EGF stimulation followed by HNRNPC irCLIP-RNP. Orange dots: EGF-responsive RDAPs. **(d)** Bubble plot indicates -$\log_{10}$(P-values) of top GO terms for cellular components, biological processes, and molecular functions of the 19 EGF-responsive RDAPs. **(e)** STRING network of HNRNPC and the 19 EGF-responsive RDAPs. Red and blue nodes: EGF-induced or reduced, respectively. Triangle-shaped nodes: canonical interactors according to STRING. Yellow node border: RDAPs part of spliceosome according to KEGG and Reactome databases. Lines: interactions within each complex; line width: interaction confidence from text mining, databases, experiments, and co-expression. Dotted lines: edges between clusters. Protein-protein interaction (PPI) enrichment network P-value was determined by STRING. **(f)** Workflow of CAPRIN1 irCLIP-RNP of sodium-arsenite-treated HEK293T leading to stress-granule (SG) formation (n = 2). **(g)** Nitrocellulose image indicates the infrared signal of CAPRIN1 irCLIP-RNP ligations in sodium arsenite-treated HEK293T samples. **(h)** Venn diagram indicates the overlap between CAPRIN1 RDAPs and previously published proteins associated with SG formation (MSGP database[11]; RNP granule database[12]). **(i)** Scatter plot indicates the $\log_2$FC values against time point 0 of differentially enriched RDAPs during SG formation followed by CAPRIN1 irCLIP-

RNP. Orange dots: SG-responsive RDAPs. **(j)** Heatmap indicates $\log_2$FC values against time point 0 calculated from imputed $\log_2$-transformed LFQ intensities for the 11 SG-responsive RDAPs. *: SG-responsive RDAP. Clusters were determined using Euclidean distance and “complete” clustering methods.

**Extended Data Figure 7: Splicing analysis of HNRNPC and RDAP-depleted A431 cells. (a)** Workflow of splicing analysis after the knockdown (KD) of HNRNPC and RDAPs (CSDE1 and UPF1) in EGF-stimulated (1h) A431 cells (n = 2). **(b)** Bars indicate knockdown (KD) efficiencies of HNRNPC and RDAPs in EGF-stimulated HNRNPC and RDAPs KD samples, as detected by qPCR. Data are represented as mean (n = 2). **(c)** Heatmap indicates Z score of normalized counts for CSDE1, UPF1, and HNRNPC in EGF-stimulated HNRNPC and RDAPs KD samples. **(d)** Stacked bars indicate the percentage of significant AS events compared to control siRNAs in EGF-stimulated HNRNPC and RDAPs KD samples. **(e)** Bars indicate the overlap of significant AS events between HNRNPC and RDAP knockdown samples after EGF stimulation. **(f)** Representative immunostaining images for HNRNPC (green) and UPF1 (red) in A431 cells after HNRNPC and UPF1 KDs. DAPI (blue signal) was used to outline the nuclear shape. Scale bars represent 20 µm. **(g)** Bars indicate the fold change against control siRNA of HNRNPC and UPF1 fluorescence intensity in HNRNPC and UPF1 knockdown A431 cells. **(h)** Representative images of proximity ligation assay (PLA) for HNRNPC and UPF1 in A431 cells after HNRNPC and UPF1 KDs. DAPI (blue signal) was used to outline the nuclear shape. Scale bars represent 20 µm. **(i)** Bars indicate the fold change against control siRNA calculated from the number of HNRNPC and UPF1 PLA foci quantified in HNRNPC and UPF1 knockdown A431 cells. **(j)** Immunoblot images indicate phosphorylated ERK, HNRNPC, and UPF1 protein signals in EGF-stimulated (0, 15, 30, 60min) A431 cells. **(k)** Immunoblot images indicate protein signal of HNRNPC and UPF1 as well as of the nuclear marker Lamin A/C and cytoplasmic marker Tubulin after subcellular fractionation of A431 cells. **(l)** Stacked bars indicate the percentage of HNRNPC and UPF1 protein signals in nuclear and cytoplasmic fractions. Data are mean ± s.d (n = 3). In **g** and **i**, data are mean ± s.d; *P-value < 0.05, **P-value < 0.01, ***P-value < 0.001 – one-way ANOVA with Tukey’s test against control siRNA cells (n = 3).

**Extended Data Figure 8: UPF1 and HNRNPC regulate the stability of intro-retained RNA transcripts during EGF stimulation. (a)** Bars indicate the overlap of significant RI events between EGF-stimulated HNRNPC-KD and UPF1-KD samples. 343 common RI events (280 genes) with congruent $\Delta$PSI patterns after HNRNPC-KD and UPF1-KD were selected. **(b)** Bubble plot indicate -log$_{10}$(P-value) of top GO terms for molecular functions, cellular components, regulatory processes, and pathways terms (KEGG, Reactome, WikiPathways) of 280 common HNRNPC-KD/UPF-KD RI genes. **(c)** Workflow of differential transcript expression analysis of EGF-stimulated HNRNPC- and UPF1-KD data compared to control siRNAs. 186 RI transcripts were found to be commonly differentially expressed (DE, q-value $< 0.05$) in EGF-treated HNRNPC and UPF1 KD samples. **(d)** Scatter plots indicate log$_2$FC values against control siRNA of 186 common DE transcripts in EGF-stimulated HNRNPC-KD and UPF1-KD samples. Red and black lines were fitted to the points using robust linear regression. Statistical comparison of regressions between common HNRNPC-KD/UPF1-KD RI transcripts and background transcripts was made using Chow's test. Dark red dots: 186 common DE RI transcripts; black dots: significant DE background transcripts. **(e)** Lines indicate the cumulative fraction of log$_2$FC values against control siRNA of 186 common DE transcripts in EGF-stimulated HNRNPC-KD and UPF1-KD samples. Red lines: 186 DE RI transcripts; black and grey lines: significant DE background transcripts in HNRNPC and UPF1 knockdown samples. P-values between common HNRNPC-KD/UPF1 RI transcripts and significant DE background transcripts were calculated by performing the Kolmogorov-Smirnov test. **(f)** Workflow of mRNA stability analysis using actinomycin D after the knockdown of HNRNPC and UPF1 in EGF-stimulated (1h) A431 cells. 158 RI transcripts were found to be commonly differentially stable (DS) after actinomycin D in EGF-treated HNRNPC and UPF1 KD samples. **(g)** Bars indicate knockdown efficiencies of HNRNPC and UPF1 in HNRNPC and UPF1 knockdown A431 cells treated with EGF for 1h and then with actinomycin D for a maximum of 4h, as detected by qPCR. Data are mean ± s.d (n = 2). **(h)** mRNA stability profiles for HNRNPC and UPF1 knocked down transcripts as a quality check of actinomycin D treatment, determined by qPCR, in HNRNPC and UPF1 knockdown samples with EGF for 1h and then with actinomycin D for a maximum of 4h (n=2). Data are mean ± s.d. **(i)** Principal component analysis (PCA) plots of EGF-treated HNRNPC and UPF1 KD samples followed by actinomycin D time course. Green dots: 0h; blue dots: 1h; purple dots: 3h; red dots: 4h actinomycin D timepoints. **(j)** Scatter plots indicate $log_2FC$ values against control siRNA of 158 common DS transcripts in EGF-stimulated HNRNPC-KD and UPF1-KD samples. Red and black lines were fitted to the points using robust linear regression. Statistical comparison of regressions between common DS HNRNPC-KD/UPF1-KD RI transcripts and background transcripts was made using Chow's test. Dark red dots: 158 common DS RI transcripts; black dots: significant DE background transcripts. **(k)** Lines indicate the cumulative fraction of $log_2FC$ values against control siRNA of 158 common DS transcripts in EGF-stimulated HNRNPC-KD and UPF1-KD samples. Red lines: 158 DS RI transcripts; black and grey lines: significant DS background transcripts in HNRNPC and UPF1 knockdown samples. P-values between common HNRNPC-KD/UPF1 RI transcripts and significant DE background transcripts were calculated by performing the Kolmogorov-Smirnov test. **(l)** Venn diagram indicates the overlap between the 35 co-bound RI genes by HNRNPC and UPF1 and the genes harboring transcripts that were differential expressed or stable (DE and DS) after HNRNPC and UPF1 KD in EGF-treated A431 cells. **(m)** Schematic of the minigene construct for RND3 and RPS9 intron retained regions. **(n)** Representative image of minigene assay for RND3 (left) and RPS9 (right) intron-retained regions in EGF-treated (1h) HNRNPC and UPF1 KD samples. Unspliced and spliced regions are indicated with blue-orange-blue and blue-blue boxes, respectively. **(o)** Stacked bars indicate the percentage of unspliced and spliced minigene fragments for RND3 and RPS9 intron-retained regions in EGF-treated (1h) HNRNPC and UPF1 KD samples. Data are mean ± s.d; **P-value < 0.01, ***P-value < 0.001 – one-way ANOVA with Tukey's test against control siRNA cells (n = 3).

**Extended Data Figure 9: UPF1 supports HNRNPC RNA splicing surveillance of EGF-regulated proliferation mRNAs. (a)** Workflow of HNRNPC and UPF1 irCLIPv2 as well as HNRNPC-UPF1 Re-CLIP in EGF-stimulated (0, 15, 30, and 60min) A431 cells (n = 2). **(b)** Nitrocellulose images indicate the infrared signal of HNRNPC and UPF1 irCLIPv2 ligations in EGF-stimulated (0, 15, 30, and 60min) A431 cells. **(c)** Nitrocellulose image indicates the infrared signal of HNRNPC-UPF1 Re-CLIP ligations from EGF-stimulated (0, 30, and 60min) A431 cells. The signal represents pooled libraries. **(d)** Pie chart indicates the percentage of 280 common RI genes categorized as protein-coding (orange), lncRNA (yellow), or pseudogene (green). **(e)** Pie chart indicates the percentage of the 307 common RI regions residing in protein-coding genes categorized as 5’UTR (orange), 3’UTR (yellow), or CDS (green). **(f)** Workflow of HNRNPC and UPF1 irCLIPv2 after UPF1 and HNRNPC KD in EGF-stimulated (0, 60min) A431 cells (n = 2). **(g)** Nitrocellulose images indicate the infrared signal of HNRNPC and UPF1 irCLIPv2 ligations after UPF1-KD or HNRNPC-KD, respectively. **(h)** Barplots showing KD efficiencies of HNRNPC and UPF1 in EGF-stimulated HNRNPC- and UPF1-KD irCLIPv2 samples, as detected by qPCR. Data are represented as mean (n = 2). **(i)** Heatmap indicates the Z score values estimated against control siRNA in HNRNPC and UPF1 irCLIPv2 knockdown data of the 30 EGF-responsive 3’UTRs of RI transcripts co-BR by HNRNPC and UPF1. Shades of blue boxes: EGF stimulation time points. **(j)** De-novo motifs discovered in EGF-responsive co-bound and co-regulated (co-BR) 3’UTRs of RI transcripts. **(k)** Volcano plot indicates the -$\log_{10}$(FDR) and max($\log_{2}$FC) values of DE genes in bulk RNA-seq from EGF-treated (0, 15, 30, and 60min) A431 cells. Orange and blue dots: up and downregulated genes. **(l)** Heatmap indicates the Z score of normalized counts of 10 co-BR RI genes by HNRNPC and UPF1 showing time-dependent-response to EGF stimulation. Orange and blue boxes: up and downregulated genes **(m)** IGV tracks indicate normalized coverage of HNRNPC and UPF1 irCLIPv2 time-course/knockdown and HNRNPC-UPF1 Re-CLIP samples at 3’UTR of DDX3X. Tracks show the signal sum of 2 replicates. Shades of blue boxes: EGF stimulation time points. **(n)** *De novo* structure prediction using AlphaFold 3 and manual refinement provides a three-dimensional model of the protein-RNA multimer complex comprising of HNRNPC, UPF1, and 3’ UTR mRNA of DDX3X. **(o)** Immunoblot image indicate HNRNPC and UPF1 protein signal after native RNA pull-down of biotinylated RND3 and DDX3X 3’UTR binding regions in 1h EGF-stimulated A431 cells. **(p)** Bars indicate fold change against scramble control of HNRNPC and UPF1 protein signal after native RNA pull-down in 1h EGF-stimulated A431 cells. *P-value < 0.05; **P-value < 0.01, ***P-value > 0.001 – unpaired one-tailed Student’s t-test against scramble control (n = 3-6).

**SUPPLEMENTARY TABLES**

**Supplementary Table 1:** Differential enrichment analysis results of HNRNPC irCLIP-RNP UVC vs. no-UV comparison in HEK293T.

**Supplementary Table 2**: Differential enrichment analysis results of irCLIP-RNP with two RNase I concentrations (1U/µL and 0.02U/µL) for HNRNPA2B1, HNRNPC, and HNRNPU in HEK293T.

**Supplementary Table 3**: Differential enrichment analysis results of label-free irCLIP-RNP for 13 RBPs in HEK293T and HepG2.

**Supplementary Table 4**: Differential enrichment results and cell-type differential analysis of TMT-labeled irCLIP-RNP for 13 RBPs in HEK293T and HepG2.

**Supplementary Table 5**: Differential window analysis results of irCLIPv2 for 4 RBPs and 3 RDAPs in HEK293T cells.

**Supplementary Table 6**: Differential enrichment results of TMT-labeled irCLIP-RNP for HNRNPC after stimulation with EGF in A431 cells.

**Supplementary Table 7**: Differential enrichment results of label-free irCLIP-RNP for CAPRIN1 during stress granule formation in HEK293T cells.

**Supplementary Table 8**: Alternative splicing events defined by rMATS after the knockdown of 4 RBPs during EGF stimulation in A431 cells.

**Supplementary Table 9**: Differential transcript expression results of mRNA splicing data after knockdown of HNRNPC and UPF1 during EGF stimulation in A431 cells.

**Supplementary Table 10**: Differential transcript stability results of mRNA stability data after the knockdown of HNRNPC and UPF1 after EGF stimulation and actinomycin D treatment in A431 cells.

**Supplementary Table 11**: Differential expression analysis results of EGF time course in A431 cells.

**Supplementary Table 12**: List of antibodies, cell lines, reagents, plasmids, siRNAs, and primers used in the current study.

### Methods

#### Cell culture

HepG2 and A431 were obtained from the American Type Culture Collection (ATCC). A431 and HEK293T (Takara Bio) cells were cultured in Dulbecco's modified Eagle's medium (DMEM) media (Gibco) supplemented with 10% fetal bovine serum (FBS, Corning) and 1% penicillin–streptomycin (Gibco) at 37°C in a 5% $CO_2$ humidified incubator. HepG2 cells were cultured in Eagle's minimal essential (EMEM) media (Corning) supplemented with 10% FBS and 1% penicillin–streptomycin at 37°C in a 5% $CO_2$ humidified incubator. All cells were routinely tested for mycoplasma contamination using the MycoScope PCR Mycoplasma Detection kit (Genlantis).

#### RNA isolation, reverse transcription, and qPCR

To perform quantitative RT-PCR (qRT-PCR), total RNA was extracted using RNeasy Plus mini kit (QIAGEN), and concentration was quantified using NanoDrop ONE (Thermo Fisher). Equal amounts of total RNA (usually 1µg) were reverse transcribed using the iScript cDNA Synthesis Kit (Bio-Rad), according to the manufacturer's instructions. qRT-PCR analysis was performed using the LightCycler 480 II System (Roche) with the SYBR Green Master Mix (Thermo Fishier). Samples were run in triplicates (10ng cDNA per reaction) and normalized to levels of GAPDH for each reaction. qPCR Primers are listed in **Supplementary Table 12**.

#### Western blotting

For immunoblot analysis, proteins were quantified by Pierce BCA Protein Assay kit (ThermoFisher) following the manufacturer's instructions. 10-20 µg of cell lysates were loaded per lane on a NUPAGE 4-12% Bis-Tris SDS-PAGE gel (ThermoFisher) and transferred to a nitrocellulose membrane (Promega) at 4°C. The membrane was blocked with Intercept blocking buffer (PBS, LICOR Biosciences) at room temperature for 1 hour and incubated with primary antibody at 4°C overnight. Membranes were then washed with TBS-T and incubated with secondary goat anti-mouse and goat anti-rabbit antibodies (LI-COR Biosciences) at a dilution of 1:5,000 for 1 hour at room temperature. After incubation with secondary antibodies, membranes were washed with TBS-T and visualized using the Odyssey CLx Infrared Imaging System (LI-COR Biosciences). Image processing was performed using Licor ImageStudioLite software (LI-COR Biosciences). Antibodies are listed in **Supplementary Table 12**.

**Visualization of irCLIP ligations**

irCLIP ligations were performed as previously described[13]. Briefly, ~80% confluent HEK293T, HepG2, and A431 cells were crosslinked on ice with 254nm UVC at 0.3J/cm$^2$ using a Stratalinker 2400 (Stratagene). After lysis, sonication, and clearance, protein concentration was quantified using the Pierce BCA Protein Assay Kit (ThermoFisher). 3mg/mL no-UVC and UVC extracts were then digested with different RNAse A concentrations (1100ng/mL to 8ng/mL). RNase-digested lysates were incubated with antibody pre-conjugated Protein G dynabeads (ThermoFisher) for 1h at 4°C. After washing, on-beads 3' irCLIP adapter ligations were performed overnight. The next day, ligation reactions were removed from the beads, and beads were resuspended in 10 µL of 1x LDS sample loading buffer (ThermoFisher) supplemented with 5% Beta-Mercaptoethanol (Sigma) and heated for 15min at 75°C. Samples were resolved by SDS–PAGE using NuPAGE 4-12% Bis-Tris Gels (ThermoFisher) at 180V for 45min. Resolved RNP complexes were wet transferred to nitrocellulose at 400mA for 60min at 4°C. Nitrocellulose membranes were visualized using the Odyssey CLx Infrared Imaging System (LI-COR Biosciences). To determine the ligated RNPs per cell, irCLIP ligations resulting from 1x10$^4$ cells were dot-blotted on nitrocellulose membrane (Promega) and visualized using the Odyssey CLx Infrared Imaging System (LI-COR Biosciences). The amount of ligated RNPs per cell was finally calculated after converting the fluorescence unit into the number of ligations based on the fluorescence value of the free ligation adapter (~7,500 fluorescence units/fmol, as determined by Licor ImageStudioLite software (LI-COR Biosciences)) and dividing the resulting value by the number of lysed cells.

**irCLIP-RNP**

*UVC crosslinking and extract* preparation. No-UVC and UVC-treated cells were harvested as follows. For the EGF-stimulation timecourse, A431 cells were incubated for 16h in starvation medium (DMEM, 0.01% FBS, and 1x Pen Strep) and stimulated by the addition of recombinant human EGF (R&D) to a final concentration of 50ng/mL. For induction of stress granules in HEK293T cells, growth media was supplemented with 500µM sodium arsenite (Sigma). Growth or stimulation media was rapidly decanted, and cells were washed on the plate with excess ice-cold PBS for ~20-30s. PBS was then rapidly decanted, and plates were held vertically for ~10s to allow residual PBS to collect for final vacuum-based aspiration. Cells were then immediately placed on ice. For no-UVC negative controls, cells were lysed on the plate (0.3mL or 1mL for 10cm and 15cm plates, respectively) with the ENCODE CLIP lysis buffer (50mM Tris (pH 7.5), 1% Igepal, 0.5% Na-deoxycholate, 150mM NaCl, 5mM EDTA) and lysates collected via cell scraper. For UV crosslinking, cells were placed in a UV Stratalinker 2400 (Stratagene) equipped with 254nm lamp on 'energy setting' and exposed to 0.3J/cm$^2$ and then lysed with ENCODE CLIP lysis buffer and collected as described above. Extracts were then sonicated with a Branson digital sonicator microtip (Branson) for 10s at 10% power. Insoluble material was then pelleted at 16,000rcf for 10min at 4°C and clarified extracts were transferred to new tubes. Protein concentrations were then measured using the Pierce BCA Protein Assay kit (ThermoFisher). Post quantification, cell lysates were adjusted to 3.0mg/mL and stored at -80°C. For the species-mixing experiment, we mixed 1:1 the human and mouse lysates with opposite crosslinking conditions.

*RNase digestion*. 3.0mg/mL no-UVC and UVC extracts were first brought to 4°C and then incubated for 10min in a 37°C water bath. 100x working solutions of RNAse (A or 1) were then added, samples vortexed for 10s, and then incubated for 5min at 37°C. RNase reactions were stopped by the addition of 2 volumes of ice-cold ENCODE CLIP lysis buffer. 1.0mg/mL processed lysates were then stored at -80°C. For all irCLIP-RNP experiments, biological replicates were generated at least 2 weeks apart.

*Immunoprecipitation and adapter ligation.* All irCLIP-RNP immunoprecipitations were scaled up to generate 1pmol ligated RNPs. Antibodies were pre-conjugated overnight to Protein G dynabeads (ThermoFisher) in 1x PBS at a ratio of 7.5µL of beads per 1µg of primary antibody at a final concentration of 100µg/mL. For all immunoprecipitation with mouse-IgG, -IgG1, -2a, or 2b, an equal amount of secondary rabbit anti-mouse IgG (ThermoFisher) was added. Beads were washed once with 1mL of PBS and resuspended with 1xPBS supplemented with 0.5% BSA and 0.05% sodium azide at 0.2mg/mL and stored at 4°C for up to 4 weeks. Immunoprecipitation was performed overnight and washed at 4°C as follows. For small-scale experiments, washes were: 1.0mL 1x high stringency buffer (H-STR, 50mM Tris-HCl (pH 7.5), 5mM EDTA, 1% Triton-X 100, 1% Na-deoxycholate, 0.001% SDS, 120mM NaCl, 25mM KCl) 5min with rotation, 1x high salt buffer (H-SLT, 20mM Tris-HCl (pH 7.5), 5mM EDTA, 1% Triton-X 100, 1% Na-deoxycholate, 0.001% SDS, 1M NaCl) 5min with rotation followed by two rapid rinses of 0.5mL Low-Salt-pH (1mM Tris (pH 6.5), 10mM $MgCl_2$). For larger-scale irCLIP-RNP immunoprecipitations performed in 15mL conical, beads were washed with 10mL of H-STR and 10mL H-SLT. Beads were then rinsed 1x with 5mL of Low-Salt-pH buffer and then resuspended in 1mL of Low-Salt-pH buffer and transferred to a 1.5mL tube for 3' end repair. For all immunoprecipitations smaller or equal to 1µg of primary antibody, 3' end repair was performed in 30µL of dephosphorylation buffer (1mM Tris (pH 6.5), 10mM $MgCl_2$, 5mM DTT, 0.75ul T4 PNK (NEB)) + 6µL of PEG400 for 30min at in a thermomixer at 37°C, 15s 1500rpm, 1min 45s rest. After 30min, the thermomixer was adjusted to 16°C, and an equal volume (36µL) of 1x T4 ligation reaction (50mM Tris pH 7.5, 10mM $MgCl_2$, 1mM DTT, 1pmole pre-adenylated adapter, 16.7% PEG400, 0.75µL T4 RNA ligase 1 (NEB)) was added directly to the dephosphorylation reaction and incubated overnight, 15s 1750rpm, 1min 45s rest. For large-scale irCLIP-RNP, end repair reaction volumes were scaled up linearly. When necessary, end repair reactions for a single IP were split across 2 or more tubes if the end repair reaction volume + PEG400 volume exceeded 0.75mL. For all large-scale irCLIP-RNP 3' ligations, 20pmoles of the pre-adenylated adapter were added per 1.5mL tube in 1x T4 ligation reaction buffer + 16.7% PEG in 1:1 volume as described for the small-scale samples. Antibodies are listed in **Supplementary Table 12**. The following pre-adenylated adapter was used:

| Name | Sequence |
|---|---|
| 3' adapter | 5'-/5phos/AGATCGGAAGAGCACACGTCTGAACTCCAGTCAC/Barcode 6nt/ATCTCGTATGCCGTCTAAAAAAAAAAAAAAAAAA/iAzideN/CT/3bio/-3' |

*Processing of irCLIP-RNP ligations for in-gel tryptic digestion*. Overnight irCLIP-RNP ligations were placed on a magnet to collect Protein G dynabead-bound ligated-RNPs. Beads were resuspended in 1mL of ENCODE CLIP lysis buffer and 10µL removed for immediate QC of ligated-RNPs by standard NuPAGE Bis-Tris 4-12% SDS-PAGE (ThermoFisher), nitrocellulose transfer, and quantification of irCLIP-RNP-ligations by Odyssey CLx Infrared Imaging System (LI-COR Biosciences). The remaining beads were placed on a magnet to re-collect for 2min, followed by the removal of the wash buffer. Ligated-RNPs were then eluted by the addition of 250µL of SDS Elution Buffer (50mM Tris pH 7.5, 1% SDS, 1mM EDTA, 1mM TCEP) and 10min incubation at 65°C in a thermomixer at 1500rpm. Beads were then collected on a magnet for 2min, and eluates were transferred to new low-bind 1.5mL tubes. 500µL of IP Dilution Buffer (50mM Tris pH 7.5, 3% Triton, 150mM NaCl, 1mM EDTA) was then added to each elution and 3µL removed for overnight storage at 4°C for dot blot input control. 100µL of 3x pre-washed (3x1mL eCLIP Lysis Buffer) Dynabeads MyOne Streptavidin C1 (ThermoFisher) were added to each sample and rotated overnight at 4°C. Beads were collected on a magnet for 2min, and 3µL of the unbound fraction was removed for dot blot QC (2µL of Dynabeads MyOne Streptavidin C1 input and 2µL of unbound were dot blotted onto nitrocellulose and immediately scanned on an Odyssey CLx to verify >95% capture of ligated-RNPs). After verifying capture, the remaining unbound fraction was removed, and beads were washed with 1mL H-SLT buffer in a thermomixer for 1min at 1,500rpm, 25°C. Beads were then washed twice with 1mL of 2% SDS wash buffer (50mM Tris pH 7.5, 1mM EDTA), in a thermomixer for 1min 1,500rpm, 37°C. After the final wash, beads were pelleted for 1min at 10,000g before being placed on a magnet for 2min. All traces of the final wash were removed and ligated-RNPs eluted by the addition of 30µL of 1xLDS supplemented with 5mM-BME, and 2mM-Biotin and incubation in a thermomixer for 30min, 2,000rpm, 75°C. Samples were then stored at -20°C for in-gel tryptic digestion.

*In-gel tryptic digestion and peptide purification*. 30µL of Dynabeads MyOne Streptavidin C1 bead elutions were resolved by NuPage Bis-Tris 4-12% SDS-PAGE (ThermoFisher) for 30min at 200V with samples flanked on both sides with protein standards (Promega). Prior to fixation, gels were scanned on an Odyssey CLx Infrared Imaging System (LI-COR Biosciences) at 0.5mm offset. Gels were then rocked in fixation buffer (50:40:10 Methanol:AceticAcid:$H_2$0) for 10min at room temperature and equilibrated in 50mM Ammonium Biocarbonate (ABC) buffer for 10min at room temperature. After additional equilibration in 50mM-ABC/40mM-BME for 10min at room temperature, gels were sealed in a bag with 1 gel volume of 50mM-ABC supplemented with 40mM-BME and placed in a 55°C water bath for 30min. Gels were transferred to a new vessel containing 10 volumes of 50mM-ABC supplemented with 55mM-acrylamide and rocked for 30min in the dark. Gel sections were then excised as desired, and gels were crushed through 0.5mL tubes (containing 18-gauge hole punches) into 1.5mL low-bind tubes. Crushed gel was then shrunk with ~5 volumes of a 50:50 mix of 50mM-ABC and acetonitrile and incubation in a 25°C thermomixer for 10min, 750rpm. A P200 pipet was used to carefully remove shrink solution above the gel bed. An additional 5 volumes of 50:50 50mM-ABC:acetonitrile was added for a second round of incubation in a 25°C thermomixer for 10min, 750rpm. A P200 pipet was again used to carefully remove shrink solution above the gel bed. De-hydrated gel pieces were completely dried for 20min in a speed vac. For tryptic digestion, gel pieces were rehydrated for 30min on ice with 50µL Trypsin buffer (50mM ABC, 0.02% ProteaseMax (Promega), 400ng Trypsin/LysC (Promega). After 30min, remaining exposed gel was covered with 50mM-ABC+0.02%ProteaseMax and incubated o/n at 37°C. Peptides were then extracted by addition of 200µL of extraction buffer (acetonitrile:H20:formic-acid, 70:29:1) and 10min incubation in a thermomixer, 37°C, 15s

900rpm, 45s rest. Gel pieces were pelleted for 5s at 10,000rpm, and supernatant was transferred to a new low-bind tube. Extraction was repeated with an additional 200µL and pooled with the first extraction. Acetonitrile was then removed by 1h speed-vac followed by -80°C lyophilization to complete dryness. Dried peptides were resuspended in 400µL 0.5% Formic Acid and stored at -80°C until C18 column clean-up.

*C18 column clean up and on-column TMT labeling*. Monospin C18 Columns (GL Sciences Inc.) were equilibrated by two 200µL washes with 50:50 acetonitrile:50mM-ABC at 6,000rpm for 1min each. Columns were then rinsed twice with 200µL of 0.1% Formic acid at 6,000rpm for 1min each. All liquid was completely removed from the flow through the tube, and acidified peptides were captured on the column at 6,000rpm for 1min. Flow through was then re-loaded and spun through the column for 1min at 6,000rpm. For non-TMT labeled samples, columns were washed twice with 200µL of 0.1% Formic Acid at 6,000rpm for 1min each, and samples were eluted into a clean tube with 400µL of elution buffer (acetonitrile:H20:Formic-Acid, 60:40:0.1) at 6,000rpm for 2min. For on-column TMT-labeled peptides, 100µL of PBS was used to resuspend 20µg of 1.5ul TMT aliquots (ThermoFisher), which was then immediately added to the C18 column for 5 min. Columns were spun for 1min at 6,000rpm and then washed twice and eluted as described for non-TMT labeled peptides. Acetonitrile was then removed by 1h speed-vac followed by -80°C lyophilization to complete dryness. Dried peptides were then resuspended in 10µL 0.1% and stored at 4°C prior to mass spectrometry. LC-MS/MS experiments were conducted on a Q Exactive Plus mass spectrometer equipped with an UltiMate 3000 UPLC system (Thermo Fisher Scientific).

**irCLIP-RNP data analysis**

For all label-free irCLIP-RNP data, Maxquant (ver. 2.0.1.0) was used for protein identification and quantification in LFQ mode[14], searching against the human Uniprot database, Proteome ID UP000005640 (reviewed and with isoforms). The following settings were used: maximum number of miss-cleavages for trypsin: 2 per peptide; cysteine carbamidomethylation: fixed; methionine oxidation: variable modifications; tolerances in mass accuracy: 20 ppm for both MS and MS/MS. maximum false discovery rates (FDRs): 0.01 at both peptide and protein levels; minimum required peptide length: 6 amino acids. Before any downstream analysis, we removed contaminants and reverse proteins.

*Label-free HNRNPC irCLIP-RNP reproducibility test.* We analyzed differential enrichment using the DEP2 R package[15] following the authors' tutorial page. Briefly, we first selected proteins detected in all replicates of at least one crosslinking condition as well as the isoform with the highest intensity across samples. Then, $log_2$-transformed LFQ intensities were normalized by performing a variance stabilizing transformation using the normalize_vsn() function. Normalized LFQ intensities were imputed using the quantile regression-based left-censored function (QRILC) through the impute() function. Differential enrichment analysis was performed by comparing UVC samples from the whole RNP zone against UVC samples from the free RNA ligation zone and no-UV samples from both zones. In the test_diff function, UVC samples from the “whole RNP zone” were set as “control” contrast. Given the lower number of detected proteins, FDR was calculated on the DEP2-generated uncorrected P-values using the p.adjust() function. Proteins showing an FDR < 0.05 and an FC value > 3 against no-UV samples from the whole RNP zone were considered RDAPs. Analysis results are reported in **Supplementary Table 1**.

*Species-mixing label-free HNRNPC irCLIP-RNP.* Unique peptide intensities retrieved from the peptide.txt MaxQuant output file were first filtered for contaminants and reversed proteins. Then, unique peptide intensities were normalized and imputed using the normalize_pe() and impute_pe() functions from the DEP2 R package[15] following the authors’ guidelines. ROC analysis was performed by calculating the cumulative fraction using the ecdf() R function. Lollipop plots of unique peptide intensities across human and mouse proteins were done using the trackViewer R package[16].

*Label-free irCLIP-RNP of 3 RBPs comparing RNase concentrations*. First, we removed the proteins detected in the IgG sample and selected proteins that were detected in both replicates of at least one RNase I condition. To determine UVC-enriched proteins (a.k.a. RDAPs), we performed P-value independent FDR calculation between no-UV (1 replicate) and UVC samples (2 replicates) using the Clipper R package[17] following the authors’ tutorial. Only proteins with FDR < 0.1 and FC > 3 against no-UV samples were determined as RDAPs. To perform differential enrichment analysis, we used the DEP2 package, as described above, by comparing 1U/µL against 0.02U/µL RNase I concentrations. In the test_diff function, 0.02U/µL RNase I concentration was set as “control” contrast. RDAPs showing an FDR < 0.05 and a $log_2$FC value < 0 were selected. Analysis results are reported in **Supplementary Table 2**.

*Label-free irCLIP-RNP analysis of 13 RBPs*. High-confident RDAPs were selected from the proteins detected after MaxQuant analysis using the Clipper tool[17], as described above. Only proteins with FDR < 0.1 and FC > 3 against no-UV samples in at least one cell line and one RBP were selected. Gene ontology analysis was performed using gProfileR[18] webtool (https://biit.cs.ut.ee/gprofiler/) with default settings. GO terms with P-value < 0.05 were selected. For correlation analysis, RDAPs $log_2$-transformed LFQ intensities were first normalized using normalize_vsn() function and imputed from QRILC using the impute() function as described above. Finally, Pearson correlation was calculated using the cor() R function on imputed LFQ intensities. Clustering was performed using correlation distance and Ward clustering methods. Analysis results are reported in **Supplementary Table 3**.

*Label-free irCLIP-RNP of CAPRIN1 after sodium arsenite*. The selection of high-confident RDAPs was performed as described above. After normalizing and imputing the $log_2$-transformed LFQ intensities, differential enrichment analysis was performed against time point 0 using the limma R package[19] with the following design formula (~ sodium_arsenite_stimulation_time). Proteins with FDR < 0.1 across the treatment and an abs($log_2$FC value) > 1 in at least one time point were defined as stress granule-responsive RDAPs. Analysis results are reported in **Supplementary Table 7**.

For all TMT-labeled irCLIP-RNP data, Maxquant (ver. 1.6.17.0) was used for protein identification searching against the human Uniprot database, Proteome ID UP000005640 (reviewed) using the Emmot lab TMT-16plex workflow. Before any downstream analysis, we removed contaminants and reverse proteins.

*TMT-labeled irCLIP-RNP of 13 RBPs*. To define RDAPs, we performed a differential enrichment analysis by comparing UVC vs. no-UV samples using the DEP2 R package[15], as described above. We performed the analysis for each cell type separately. Proteins showing an FDR < 0.1 and an FC value > 1.2 against no-UV samples in at least one RBP and one cell type were defined as RDAPs. To increase the confidence, TMT-detected RDAPs were further filtered for those also defined as RDAPs in the label-free irCLIP-RNP experiment described above. To characterize the cell-type-specific multi-protein assembly tendency of RDAPs, $log_2$-transformed TMT intensities were first normalized using variance stabilizing transformation, and differential enrichment analysis was performed using the limma R package[19] and a two-factor interaction design formula (~ cell_type*RNP_subzone). Proteins with FDR < 0.05 and a $log_2$FC value > 0 or < 0 were defined as high-MW or low-MW RDAPs, respectively. Among these, proteins showing a cell-type:RNPsubzone interaction FDR < 0.1 were determined as RDAPs with cell-type-selective distribution. To visualize cell-type differences in RDAP distribution, we estimated slope values across the three RNP subzones from normalized $log_2$-transformed TMT intensities by performing linear regression analysis using the lm() R function (**Extended Data Fig. 4c**). Network analysis and community detection were performed using a spin-glass model and simulated annealing available in the igraph R package. Analysis results are reported in **Supplementary Table 4**.

*TMT-labeled irCLIP-RNP of HNRNPC after EGF*. We first filtered the RDAPs by performing differential enrichment analysis between UVC and no-UV samples of HNRNPC irCLIP-RNP generated from untreated A431 cells using the Clipper tool[17], as described above. Only proteins with an FDR < 0.1 and a FC > 3 were considered RDAPs. This is because any no-UV samples were run for this dataset, given the limited size of the TMTPro 16plex kit (ThermoFisher). To identify RDAPs responsive to EGF stimulation, $log_2$-transformed TMT intensities were first normalized as described above, and differential enrichment analysis was performed against time point 0 using the limma R package[19] and a two-factor interaction design formula (~ RNP_subzone + RNPsubzone:EGF_stimulation_time). RDAPs with FDR < 0.1 across the stimulation and an abs($log_2$FC value) > 0.3 in at least one time point were defined as EGF-responsive RDAPs. Gene ontology analysis was performed as described above. Protein-protein interaction network for the proteins identified after HNRNPC irCLIP-RNP-TMT after EGF stimulation was generated using the STRING webtool (https://string-db.org/cgi/input.pl). The human PPI database was used for the analysis, and interaction confidence was defined from text mining, databases, experiments, and co-expression. MCL clustering was performed using an inflation rate value of 3. Analysis results are reported in **Supplementary Table 6**.

**Re-CLIP**

After performing immunoprecipitation with the first RBP and RNA ligation as described above, irCLIP-RNP ligations were resuspended in 100µL Nuclei Lysis Buffer (50mM Tris pH 7.5, 1%SDS, 1mM EDTA, 250µM TCEP) and incubated for 10min at 65°C with shaking (1500rpm). Eluted ligated RNPs were then transferred to a new tube containing 4 volumes of ice-cold Triton Dilution Buffer (50mM Tris pH 7.5, 125mM NaCl, 2.5% Triton-X). Samples were pre-cleared with the same amount of Protein G dynabeads (ThermoFisher) as the first immunoprecipitation for 2h at 4°C. Immunoprecipitation was performed overnight at 4°C with Protein G dynabeads previously incubated overnight with the second antibody using a similar ratio of magnetic beads/antibody as for the first immunoprecipitation. Beads were washed twice with H-STR buffer, resuspended in 20µL of 1x LDS buffer with 5mM BME, and incubated for 10min at 65°C. Finally, Re-CLIP ligations were resolved on SDS-PAGE, transferred to nitrocellulose membrane, and subjected to irCLIPv2 library preparation.

**irCLIPv2 library preparation**

After overnight immunoprecipitation of irCLIP-RNP or RE-CLIP ligations performed as described above, beads were washed once with 1.0mL ENCODE CLIP lysis buffer, resuspended in 1xLDS sample buffer supplemented with 5% beta-mercaptoethanol, and heated for 15min at 75°C. Ligated RNPs were then separated by electrophoresis using NUPAGE 4-12% Bis-Tris gels (ThermoFisher) and transferred to nitrocellulose membrane (Promega). Nitrocellulose-transferred ligated RNPs were carefully excised on damp Whatman filter paper, sliced into small pieces, and transferred to a 1.5mL siliconized tube. Ligated RNPs were then recovered by the addition of 150µL of Proteinase K (PK) reaction buffer (100mM Tris pH 7.5, 100mM NaCl, 1mM EDTA, 0.5% SDS, 5ul 20/mg/mL Proteinase K (ThermoFisher)) and incubation in a 55°C thermomixer, 15s 900rpm, 45s rest. Tubes were then briefly centrifuged, and the PK reaction was transferred to a new 1.5mL tube. Nitrocellulose pieces were then rinsed with 300µL of ice-cold Biotin IP buffer (50mM Tris pH 7.5, 500mM NaCl, 2% Triton) and then combined with the initial PK elution. 7.5µL of twice washed (0.2mL Biotin IP buffer) Dynabeads MyOneC1 Streptavidin magnetic beads (ThermoFisher) were added to each sample and rotated at 4°C for 30min. Beads were placed on a magnet, and unbound material was removed. Beads were then resuspended in 100µL of H-STR buffer and transferred to a PCR tube in 8-strip format. All further bead resuspensions were done by vortexing. Beads were then rinsed 3 more times, 1x 0.1mL H-STR, 2x 0.1mL PBS. Beads were resuspended in 1x cDNA annealing mix (4µL 5x Superscript IV buffer, 1µL of 1µM P7-end cDNA primer, 15µL water) and rotated at 55°C for 10min followed by incubation for 10min at 25°C. 10µL of cDNA Reaction Mix (2µL 5X Superscript IV buffer (ThermoFisher), 0.3µL 100mM DTT, 0.2µL Superscript IV (ThermoFisher), 1.5µL 10mM dNTPs, 6µL H20) was then added, mixed by vortexing, and rotated at 25°C for 5min and then 50°C for 15min. Samples were then cooled for 10min at 25°C. 2µL of RNAse H (Enzymatics) was added, and samples were rotated for 15min at 30°C. Beads were then resuspended in 200µL PBS and split between two tubes. Each half of the beads was then resuspended in 30µL of A-tailing or C-tailing mix (3µL 10x Terminal Transferase Buffer (NEB), 1.5µL 10mM d/ddNTP, 0.25µL Terminal Transferase (NEB), 25µL water, 10mM dATP/ddATP or dCTP/ddCTP are formulated at 30:1) and rotated at 37°C for 30min. Beads were then washed 2x0.1mL PBS and eluted in 20µL water in a 98°C thermomixer, 90s rest, 30s 2,000rpm. Beads were rapidly pelleted for 5s in a PCR minicentrifuge and eluates transferred to a new PCR tube and incubated at 25°C for 5min. 5µL of 5x Phusion HF Buffer (ThermoFisher) and 1µL of Annealed-barcoded-2$^{nd}$-strand oligo were then added to each sample and incubated at 45°C for 5min followed by 0.1degree/s ramping to 25°C. 4µL of 2$^{nd}$ Strand Synthesis Reaction mix (1µL 5x Phusion HF Buffer (ThermoFisher), 1.5µL 10mM dNTPs, 0.2µL Bsu polymerase (NEB), 1.8µL water) was added and samples incubated at 25°C for 5min followed by 37°C for 15min. cDNA was purified by the addition of 2 volumes of AMPure XP beads (Beckman), incubation for 10min, 2x0.1mL 80% ethanol washes, 8min drying, and elution in 20µL water. 20µL of PCR-A (for A-tailed cDNA) or PCR-C (for C-tailed cDNA) was then added (20µL 2xPhusion HF mix, 0.25X SYBR Green I (ThermoFisher), 400nM P5-PCR1(A or C) and P7-end cDNA primer) and samples amplified as follows: 98°C 1min, 6-13 cycles (98°C 5s, 67°C 45s). PCRs were then purified by the addition of 72ul of AMPure XP beads as previously described, and DNA was eluted with 10µL water. 10µL of 2x Novex TBE-Urea sample buffer (ThermoFisher) was then added, and samples were incubated at 75°C for 10min. PCR1s were then resolved by Novex 6% TBE-UREA gel at 300V for 25min and visualized by SYBR Gold (ThermoFisher) staining. PCR amplicons greater than 20nt were then targeted for isolation (no insert = 130nt) by excision of the 150-225nt gel region. Crushed gels were then incubated overnight in 300µL of Elution Buffer (500mM NaCl, 1mM EDTA) in a 55°C thermomixer, 15s 900rpm, 45s rest. Gel pieces were removed with SPIN-X columns, and sample volumes were reduced to less than 100µL by speed-vac. DNA was then purified using Zymo DNA Clean and Concentrator Kit (Zymo Research) and 20µL water elution. Purified PCR1 DNA was then amplified for an additional 4 cycles with PCR2 mix (20µL 2XPhusion HF mix, 0.25X SYBR, 400nM P5-Solexa, 400mM P7-end cDNA primer). Samples were then purified with 72µL of AMPure XP beads as previously described. Samples were then quantified on NanoDrop ONE (ThermoFisher) and submitted for Bioanalyzer at the Stanford Genome Technology Center for precise quantification. Equimolar of A and C libraries were submitted for NGS at Novogene. The following oligos were used:

| Name | Sequence |
|---|---|
| P7-end cDNA primer | 5'-CAAGCAGAAGACGGCATACGAGAT-3' |
| Annealed-barcoded-2$^{nd}$-strand forward | 5'-TCCCTACACGACGCTCTTCCGATCT/barcode 5nt/NNNNNNNNTTTTTTTTTTTTVN-3' |

| | |
|---|---|
| Annealed-barcoded-2$^{nd}$-strand forward | 5'-TCCCTACACGACGCTCTTCCGATCT/barcode 5nt/NNNNNNNNGGGGGGGHN-3' |
| Annealed-barcoded-2$^{nd}$-strand reverse | 5'-TCAGTAGATCGGAAGAGCGT/3ddC/-3' |
| P5-PCR1 A | 5'-CGAGATCTACACTCTTTCCCTACACGACGCTCTTCCGATCT-3' |
| P5-PCR1 C | 5'-CCACCGAGATCTACACTCTTTCCCTACACGACGCTCTTCCGATCT-3' |
| P5.Solexa | 5'-AATGATACGGCGACCACCGAGATCTACACTCTTTCCCTACACGACGCTCTTCCGATCT-3' |

**RE-CLIP and irCLIPv2 data analysis**

For every Re-CLIP and irCLIPv2 sequencing dataset, pooled libraries were initially demultiplexed using the i5/i7 Illumina indexes. Raw reads were further demultiplexed into sample libraries using the tailing nucleotide sequence and the 5' barcode, while simultaneously trimming these base pairs, using Cutadapt[20] (ver. 1.8.1). Unique molecular identifiers (UMIs) were extracted using UMItools[21] (ver. 1.0). Finally, the 3' adapter was trimmed using Cutadapt, filtering out the remaining sequences under 16 bps. These libraries were aligned to repetitive RNA sequences, and unmapped reads were then aligned to the human reference genome (gencode release v39 basic annotation) using the STAR aligner (ver. 2.7.8a). These alignments were merged and sorted using samtools, and reads with identical alignments containing the same UMI were collapsed using UMItools. Samples with different nucleotide tails were merged for downstream analyses. Strand-splitted bigwig files were generated from crosslink sites extracted from merged bam files using a custom python script followed by genomecov and bedGraphToBigWig tools.

*HNRNPC Re-CLIP of ELAVL1, KHSRP, and HNRNPM*. Bigwig files were normalized as described above using the size factors estimated from the crosslink counts by the DEWSeq R package . A coverage heatmap from normalized bigwig files around HNRNPC binding regions identified in the irCLIPv2 dataset was generated using the EnrichedHeatmap[28] R package (ver. 1.26.0) following the authors' tutorial. Tracks of replicate-summed normalized bigwig files were generated using the IGV software[29].

*HNRNPC Re-CLIP of UPF1 during EGF stimulation*. Bigwig files were generated and normalized as described above. Density heatmap of Re-CLIP RT stop locations was generated as described above. Coverage profiles were determined across the 3'UTR RI regions using the normalizeToMatrix() function of the EnrichedHeatmap package. De-novo motif analysis was performed using the runStreme() function of the memes R package with default settings. Track were generated with normalized bigwigs with the IGV software as described above.

*irCLIPv2 of 4RBPs and 3 RDAPs from 3RNP subzones*. For DEWSeq analysis, merged bam files were processed with 50bp window size and 20bp window step to preprocessed annotation (gencode release v39 basic annotation), and crosslink sites were extracted and counted using htseq-clip tools[22] (ver. 1.0). Detection of enriched regions for each of the three RNP subzones over the no-UVC sample was performed using the sliding-window approach from the DEWSeq R package[23] (ver. 1.10.0) following the authors' suggested pipeline. Only windows showing an FDR $< 0.05$ and $\log_2 FC > 1$ and against no-UV samples were collapsed into binding regions using the extractRegions() function. For 3' shift analysis, coverage profiles from normalized bigwig files were generated around (+/- 50bp) the highest crosslink nucleotide inside the significant window with the highest $\log_2$FC value against no-UVC in the RNP subzone #1. Normalization of bigwig files was performed by dividing the unscaled bigwigs by the size factor retrieved from the DEWSeq analysis. For region categorization across RNP subzones, normalized counts retrieved from DEWSeq analysis were used to estimate the slope values by performing linear regression analysis using the lm() function in R. Regions were then assigned to the respective RNP subzone type after applying a ±1 standard deviation cut-off on the slope values across all the RBP tested. Distribution of categorized binding regions in the different genomic features (5'UTR, CDS, 3'UTR, intron, other exons) was done using the ChIPpeakAnno [24] R Package (ver. 3.30.1) with the gencode release v39 basic annotation following the authors' user's guide. For motif analysis, categorized binding regions were first resized to 250, 150, and 75bp downstream of the significant window with the highest $\log_2$FC value against no-UVC in the RNP subzone #1. Motif enrichment analysis was then performed using runAme() function from the memes R package[25] HNRNPC-KHSRP region pairs were determined after applying distance cut-offs of 75bp, 150bp, or 250bp from the window with the highest $\log_2$FC value against no-UVC in the respective slope category. Analysis results are reported in **Supplementary Table 5**.

*irCLIPv2 of HNRNPC and UPF1 during EGF stimulation*. Bigwig files were generated and normalized as described above. To analyze the density of irCLIPv2 signal, RT stop locations for each genomic feature (5'UTR, exon, intron, 3'UTR) were determined using the cliProfiler R package. Then, density heatmap of RT stop locations were generated using the densityHeatmap() function from ComplexHeatmap R package[26]. To analyze changes in the total signal distribution within the 3'UTR regions of RI transcripts during EGF stimulation, unscaled bigwig files were first summed at the 3’UTR RI regions using the getCountsByRegions() function from the BRGenomics R package. Next, using the DESeq2 R package[27], Z-scores to compare each time point against the initial time point 0 were calculated by first normalizing the total signal with the estimateSizeFactors() function, obtaining $\log_2$FC and standard error coefficients through the DEseq() function, and finally dividing the $\log_2$FC by the standard error to derive the Z-score. Regression analysis of Z-scores was performed by fitting a linear regression to the data using the lm() function.

**Protein-RNA structural modeling**

A modeling technique which combined *de novo* structure prediction from AlphaFold 3 (AF3) [30] and manual refinement[31] was used to develop a three-dimensional structure of the protein-RNA multimer complex comprising of HNRNPC, UPF1 and 3’ UTR mRNA of RND3 based on CLIP data. To begin the structural modeling, the canonical protein sequences of HNRNPC (UniProt: P07910) and UPF1 (UniProt: Q92900) were used. Additionally, to constrain the challenge of *de novo* structure prediction and to improve the accuracy, the protein sequences were reduced to only those sub-sequences or domains that were annotated to be interacting with RNA. Thus, for HNRNPC the residues from 1–100 and for UPF1 the residues 101–950 were selected as input for AF3. Similarly, based on CLIP data, only an 80 and 40 nucleotide RNA sequences comprising the 3’ UTR of RND3 and DDX3X were used as input for AF3, respectively. The seed used for the prediction was 1234567, and only the default five predicted models were used for further screening. The five models were visualized and screened using UCSF ChimeraX[32], of which a model with concordance with CLIP data and previous annotation was chosen as the representative structure.

**RNA sizing of irCLIP-RNP ligations**

After proteinase K digestion of nitrocellulose-transferred HNRNPC irCLIP-RNP ligations derived from three RNP subzones (#1: 60-120kDa, #2: 120-225kDa, and 3#: 225-350kDa) RNA was captured with 100µL of 3x pre-washed (3x1mL eCLIP Lysis Buffer) Dynabeads MyOne Streptavidin C1 (ThermoFisher) and rotated overnight at 4°C. RNA was then eluted in 1x RNA loading buffer (95% formamide, 1mM EDTA, 2mM Biotin, 0.02% bromophenol blue) for 10min at 75°C. RNA length was assessed after separation by electrophoresis using Novex 6% TBE-UREA gels (Thermo).

**siRNA knockdown**

To knock down a gene of interest, 0.75-1x$10^6$ A431 cells were reverse transfected in 10cm dishes with 25nM of siRNA smartpool (Horizon) and 25µL Lipofectamine RNAiMAX (ThermoFisher) previously mixed in 1.25mL Opti-MEM (Gibco). Knockdown efficiency for each siRNA was checked by qPCR using the primers listed above. The siRNA-containing media was left on the cells for 36-48h before performing downstream experiments. siRNAs are listed in **Supplementary Table 12**.

**RNA-seq**

*mRNA splicing* analysis. 1x$10^5$ A431 cells previously transfected with siRNAs were split into 6-well plates and cultured overnight. The next day, cells were starved for 8h with starvation medium and stimulated with 50ng/mL recombinant human EGF (R&D). After 60min stimulation, cells were washed once with PBS, and 350µL RLT buffer was added to the cells. After 5min incubation at room temperature, lysed cells were collected and transferred to a 1.5mL tube.

*mRNA stability analysis*. 5x$10^4$ A431 cells previously transfected with siRNAs were split into 12-well plates and cultured overnight. The next day, cells were starved for 8h with starvation medium and stimulated with 50ng/mL recombinant human EGF (R&D). After 60min stimulation, medium was changed with starvation medium containing 5µg/mL actinomycin D (Sigma). Time points 0h, 1h, 3h, and 4h were analyzed. Cells were washed once with PBS, and 200µL RLT buffer was added to the cells. After 5min incubation at room temperature, lysed cells were collected and transferred to a 1.5mL tube.

Total RNA was extracted using the RNeasy Plus mini kit (QIAGEN) according to the manufacturer’s instructions. Libraries were prepared at Novogene. Briefly, mRNA was purified from total RNA using poly-T oligo-attached magnetic beads. After fragmentation, cDNA synthesis was performed with random hexamer primers. Subsequently, libraries were subjected to end repair, A-tailing, adapter ligation, size selection, amplification, and purification. Libraries were sequenced on a NovaSeq 6000 PE150, paired-end, 150 bp length, with ~30 M reads for each sample.

*EGF-stimulation time course analysis*. A431 cells were treated for specified time points (0, 15, 30, and 60 min) with 50ng/mL recombinant human EGF, as described above. Cells were washed once and scraped directly into RLT plus buffer supplemented with 2-mercaptoethanol and passed through QIAshredder (Qiagen) column. RNA was then extracted using the RNeasy Plus mini kit (Qiagen), and libraries were prepared with 500ng of input to the QuantSeq 30 mRNA-Seq Library Prep Kit FWD for Illumina (Lexogen) following the manufacturer's protocol. Libraries were sequenced on a NovaSeq X+, paired-end, 150 bp length, with ~30 M reads for each sample.

**RNA-seq data analysis**

*Differential event-based splicing analysis.* ENSEMBL annotations (ver. 104) were used to define possible splicing events by rMATS turbo (ver. 4.1.1) according to the authors' suggested pipeline[33]. Statistics were calculated by rMATS without the "--paired" option. Only alternative splicing events with an FDR < 0.05 were selected for further analysis. GO analysis was performed with gProfileR[18] as described above. Analysis results are listed in **Supplementary Table 8**.

*Differential transcript expression (DTE) analysis*. Raw reads were quantified with Salmon[34] (ver. 0.12.0) using the gencode release v39 basic annotation with the following settings: -l A -p 8 --gcBias --numGibbsSamples 20 --validateMappings. After quantification, raw counts were imported using tximeta (ver. 1.14.1), and differential transcript expression analysis accounting for inferential uncertainty was performed using Swish[35] from the fishpond R package (ver. 2.2.0) according to the authors' tutorial. Swish function was performed with the setting x = "conditions", where conditions were control and knockdown terms. Transcripts with a q-value < 0.05 were selected. Differential transcript expression results are listed in **Supplementary Table 9**.

*mRNA decay analysis*. Raw reads were quantified with Salmon[34] and imported using tximeta, as described above. The profile of the resulting transcripts during actinomycin D treatment was statistically tested between siRNA control and HNRNPC and UPF1 knockdown samples by performing differential transcript expression analysis accounting for inferential uncertainty using the Swish tool[35] from fishpond and limma[19] R packages. A two-factor interaction design formula (~ ActinomycinD_stimulation_time + ActinomycinD_stimulation_time:knockdown_condition) was used to analyze inferencial replicate counts previously normalized with the scaleInfReps() function. Only Transcripts with an FDR < 0.05 were selected as transcripts where their stability is influenced by HNRNPC or UPF1. mRNA stability results are listed in **Supplementary Table 10**.

*EGF-stimulation time course analysis*. Raw reads were quantified with Salmon[34] as described above. Gene-level analysis was performed using the R package DESeq2 following the authors' tutorial. Only genes with an FDR < 0.05 and an abs(max($\log_2$FC)) >

1 across EGF stimulation were selected as EGF-responsive genes. Analysis results are listed in **Supplementary Table 11**.

**Proximity Ligation Assay (PLA)**

To analyze protein-protein interaction in living cells, $5x10^4$ A431 cells were distributed in 8-well glass slide Lab-Tek II Chamber Slide (Thermo Fisher) dishes and cultured overnight. The next day, cells were starved with starvation medium. After 8h, starvation medium was replaced with starvation medium containing 50ng/mL recombinant human EGF (R&D). Time points 0min, 30min, and 60min were analyzed. Cells were then fixed with 4% PFA (ThermoFisher) diluted in DPBS (Gibco) for 10min at room temperature. Cells were washed three times with PBS and protein-protein interaction in living cells was measured by PLA with Duolink In Situ Orange Starter Kit Mouse/Rabbit (Sigma) according to the manufacturer's instructions. Anti-HA (CST, 1:200 dilution) and anti-HNRNPC (Santa Cruz Biotechnology, 1:200 dilution) antibodies were applied for PLA primary antibody incubation. After Duolink PLA probe incubation, ligation, and amplification, PLA samples were imaged by Zeiss LSM880 inverted confocal microscopy (Stanford Cell Sciences Imaging Facility), and Z-stacks of fluorescence images spanning over the entire cell monolayer were acquired. A self-built macro developed with ImageJ[36] was used to quantify the PLA foci and to count nuclear shapes (DAPI signal) on max intensity Z projections. The validity of the primary antibodies was checked by immunostaining and PLA of untreated A431 cells after the knockdown of HNRNPC and UPF1 performed as described above. For immunostaining, cells were fixed with 4% PFA (ThermoFisher) for 10min at room temperature, blocked with 5%BSA (Sigma) in DPBS (Gibco) for 1h at room temperature, and incubated overnight at 4°C with primary antibodies diluted in 2% BSA in DPBS. The next day, cells were incubated with 488 and 555-AlexaFluor anti-mouse and anti-rabbit (ThermoFisher) antibodies at 1:5000 dilution in 2%BSA in DPBS for 1h at room temperature and, finally, mounted and imaged as described above.

**Native biotinylated RNA pull-down**

*RI region cloning*. genomic DNA was isolated from around $2x10^6$ A431 cells using the DNeasy Blood and Tissue Kit (Qiagen). KHSRP-HNRNPC co-bound regions in HNRNPC and PTBP2 genes, as well as 3'UTR regions for RND3 and DDX3X, were PCR amplified from A431 genomic DNA, using PrimeStar Max DNA polymerase (Takara), according to the manufacturer's protocol. Amplified fragments were cloned into linearized pGeneArt1 (pGA.1, ThermoFisher) vector using NEBuilder HiFi DNA Assembly Master Mix (NEB) for

1h at 50°C. Ligated vectors were transformed into Stellar Competent Cells (Takara), following the manufacturer's instruction. Inserted RI regions sequence was checked by Primordium Labs. Two G-block-synthesized scramble controls were also cloned into pGA.1 vector. The amplification and cloning primers as well as scramble control sequences are listed in **Supplementary Table 12**.

*In vitro transcription and RNA biotinylation*. 1µg linearized RI-region-containing pGA.1 vectors was used as a template in an in vitro transcription reaction for 2h at 37°C, using the MEGAscript T7 transcription kit (ThermoFisher) following the manufacturer's instruction. In vitro transcription reaction was diluted with 50µL RNase-free water, and in-vitro transcribed RNA was then purified using Spin Columns and Elution Tubes (ThermoFisher). After determining the concentration, the RNA integrity was tested by agarose-formaldehyde gel electrophoresis. RNA labeling was performed using the Pierce RNA 3' End Desthiobiotinylation kit (ThermoFisher) following the manufacturer's instructions.

*Cell lysate preparation*. $6x10^6$ A431 cells per replicate were distributed in 15cm dishes and cultured overnight. The next day, A431 cells were starved and stimulated with recombinant human EGF (R&D) as described above. After 1h EGF treatment, cells were washed once with DPBS and were lysed in 600µL Pierce IP lysis buffer (ThermoFisher) supplemented with Protease Inhibitor Cocktail (Sigma). Lysed cells were incubated for 10min on ice and centrifuged at 16,000g for 5min at 4°C. Cleared lysate was transferred into a fresh tube. Protein concentration was determined by Pierce BCA Protein Assay kit (ThermoFisher).

*RNA pull-down*. Pull-down was performed using the Pierce Magnetic RNA-Protein Pull-down kit (ThermoFisher) according to the manufacturer's instructions. Briefly, 50µL per sample of streptavidin magnetic beads were washed twice with 20mM Tris (pH 7.5). After the final wash, magnetic beads were resuspended in 50µL RNA capture buffer (20mM Tris (pH 7.5), 1M NaCl, 1mM EDTA). 50pmol of biotinylated RNA was added to the resuspended magnetic beads and incubated at 25°C for 1h. RNA-bound magnetic beads were washed twice with 20mM Tris (pH 7.5) and incubated with 200µg lysate in 200µL RNA-protein binding buffer (20mM Tris (pH 7.5), 50mM NaCl, 2mM $MgCl_2$, 0.1% Tween-20, 15% Glycerol) for 2h at 4°C. Magnetic beads were carefully washed five times with RNA Wash Buffer (20mM Tris (pH 7.5), 10mM NaCl, 0.1% Tweem-20). Bound proteins were finally eluted with 1.6x LDS sample loading buffer (ThermoFisher) supplemented with 5% Beta-Mercaptoethanol (Sigma) for 15min at 75°C. Western blotting of RNA pull-down samples using HNRNPC (Santa Cruz Biotechnology, 1:500 dilution), KHSRP (Santa Cruz

Biotechnology, 1:500 dilution), and UPF1 antibodies (CST, 1:1,000 dilution) was performed as described above.

**Subcellular fractionation**

A431 cells were trypsinized and washed with PBS. Pellets were split equally into two tubes. Pellet A was lysed with 5 volumes of RIPA buffer (25 mM Tris HCl pH 7.6, 150 mM NaCl, 1% NP-40, 1% sodium deoxycholate, 0.1% SDS, 1x protease inhibitor cocktail) representing the whole cell lysate. Pellet B was resuspended in 2.5 volumes of Buffer A (10mM HEPES, 1.5mM MgCl2, 10mM KCl, 1mM DTT, 1x protease inhibitor cocktail) and incubated on ice for 1-2 minutes. An equal volume of Buffer A supplemented with 0.4% NP-40 was added to the sample and mixed by pipetting. Samples were then incubated on ice for another 5 minutes and spun down for 1 min at 4000rpm. Supernatants were taken for cytoplasmic fraction, and nuclear pellets were resuspended in 5 volumes RIPA Buffer (25 mM Tris HCl pH 7.6, 150 mM NaCl, 1% NP-40, 1% sodium deoxycholate, 0.1% SDS, 1x protease inhibitor cocktail). Nuclear lysates were incubated on ice for 30 minutes, with occasional vortexing, and then passed through a 27g needle. All lysates were cleared with centrifugation at max speed for 10 minutes, and then supernatants were quantified by BCA. Successful fractionation was confirmed, and then equal volumes were loaded for whole-cell lysate, cytoplasmic, and nuclear fractions. Three independent replicates were performed.

**Minigene assay**

*Cloning RI regions*. full-length RI regions of RND3 and RPS9 were PCR amplified from A431 genomic DNA isolated as described above using PrimeStar Max DNA polymerase (Takara), according to the manufacturer’s protocol. Amplified fragments were cloned into linearized pTAU lentivector (Addgene) using NEBuilder HiFi DNA Assembly Master Mix (NEB) for 1h at 50°C. Ligated vectors were transformed into Stellar Competent Cells (Takara), following the manufacturer’s instruction. Inserted RI region sequence was checked by Primordium Labs. The amplification and cloning primers, as well as scramble control sequences, are listed in **Supplementary Table 12**.

*Virus production*. 5x$10^6$ HEK293T cells passaged in a 10 cm plate and cultured overnight. On the following day, HEK293T cells were transfected with 7.5µg of p8.91, 3.75µg pMDG, and 7.5µg pLEX-uORF-FHH-UPF1 in 1.25mL Opti-MEM (Gibco) containing 25µL Lipofectamine 3000 (ThermoFisher). Media was replaced after 16 hours. Viral-containing media was collected 24 and 48 hours after transfection, filtered through a 0.45 um PES membrane, and concentrated to 50X with Lenti-X concentrator (Takara).

*A431 infection*. the virus titer was first determined using the Lenti-X Go-Stix Plus kit (Takara), according to the manufacturer's instructions. Then, the optimized viral titer (1 multiplicity of infection (MOI)) was added to A431 cells together with polybrene (5 µg/mL) and incubated overnight. The next day, fresh media was added, and cells were selected using puromycin (1 µg/mL) over 2 weeks before performing downstream experiments.

*Minigene amplification*. After the knockdown of HNRNPC or UPF1 using siRNAs as described above. Minigene-expressing A431 cells were lysed in RLT plus buffer and subjected to RNA isolation using the RNeasy Plus kit (Qiagen). cDNA synthesis was performed using the iScript cDNA Synthesis Kit (Bio-Rad), according to the manufacturer's instructions. Amplification of RI regions was performed using PrimeStar Max DNA polymerase (Takara), according to the manufacturer's protocol. Gel electrophoresis was performed using a 2% agarose gel stained with SYBRsafe (ThermoFisher) and imaged on a Invitrogen iBright imaging system (ThermoFisher).

**Statistical analysis of biological data**

If not alternatively specified, all results are presented as the mean with SD. Statistical tests performed to calculate P-values are specified in the related figure legend. Statistical analyses were performed with GraphPad Prism version 9 (GraphPad Software, La Jolla, CA) or R versions 4.2.0 and 4.2.1. Statistical significance was determined as follows: *P-value < 0.05, **P-value < 0.01; ***P-value < 0.001; ns, not significant.

**Data availability**

Further information and requests for resources and reagents should be directed to and will be fulfilled by the corresponding author, Paul A. Khavari. All unique/stable reagents generated in this study are available from the corresponding author without restriction. All the mass spectrometry raw and search files were deposited to the ProteomeXchange Consortium via the PRIDE[37] partner repository under the following identifier: PXD053494. All the sequencing data files are available from GEO under the following identifiers: GSE271167 and GSE271168. Codes are available on Figshare under the following links: https://figshare.com/s/283667708d45e07a2b5b and https://figshare.com/s/9d9496ff633104a600f0. Any additional information required to reanalyze the data reported in this paper is available from the corresponding author upon request.

### Supplementary information reference

1. Botti, V., McNicoll, F., Steiner, M. C., Richter, F. M., Solovyeva, A., Wegener, M., Schwich, O. D., Poser, I., Zarnack, K., Wittig, I., Neugebauer, K. M. & Müller-McNicoll, M. Cellular differentiation state modulates the mRNA export activity of SR proteins. *J. Cell Biol.* **216,** 1993–2009 (2017).
2. Avraham, R. & Yarden, Y. Feedback regulation of EGFR signalling: decision making by early and delayed loops. *Nat. Rev. Mol. Cell Biol.* **12,** 104–117 (2011).
3. Park, Y. M., Hwang, S. J., Masuda, K., Choi, K.-M., Jeong, M.-R., Nam, D.-H., Gorospe, M. & Kim, H. H. Heterogeneous nuclear ribonucleoprotein C1/C2 controls the metastatic potential of glioblastoma by regulating PDCD4. *Mol. Cell. Biol.* **32,** 4237–4244 (2012).
4. Mulnix, R. E., Pitman, R. T., Retzer, A., Bertram, C., Arasi, K., Crees, Z., Girard, J., Uppada, S. B., Stone, A. L. & Puri, N. hnRNP C1/C2 and Pur-beta proteins mediate induction of senescence by oligonucleotides homologous to the telomere overhang. *Onco Targets Ther* **7,** 23–32 (2013).
5. Mo, L., Meng, L., Huang, Z., Yi, L., Yang, N. & Li, G. An analysis of the role of HnRNP C dysregulation in cancers. *Biomark Res* **10,** 19–10 (2022).
6. Guo, W., Huai, Q., Zhang, G., Guo, L., Song, P., Xue, X., Tan, F., Xue, Q., Gao, S. & He, J. Elevated Heterogeneous Nuclear Ribonucleoprotein C Expression Correlates With Poor Prognosis in Patients With Surgically Resected Lung Adenocarcinoma. *Front Oncol* **10,** 598437 (2020).
7. Attig, J., Ruiz de Los Mozos, I., Haberman, N., Wang, Z., Emmett, W., Zarnack, K., König, J. & Ule, J. Splicing repression allows the gradual emergence of new Alu-exons in primate evolution. *Elife* **5,** (2016).
8. Wee, P. & Wang, Z. Epidermal Growth Factor Receptor Cell Proliferation Signaling Pathways. *Cancers (Basel)* **9,** 52 (2017).
9. Geiger, T., Wehner, A., Schaab, C., Cox, J. & Mann, M. Comparative proteomic analysis of eleven common cell lines reveals ubiquitous but varying expression of most proteins. *Mol Cell Proteomics* **11,** M111.014050 (2012).
10. Porter, D. F., Miao, W., Yang, X., Goda, G. A., Ji, A. L., Donohue, L. K. H., Aleman, M. M., Dominguez, D. & Khavari, P. A. easyCLIP analysis of RNA-protein interactions incorporating absolute quantification. *Nat Commun* **12,** 1569–16 (2021).
11. Nunes, C., Mestre, I., Marcelo, A., Koppenol, R., Matos, C. A. & Nóbrega, C. MSGP: the first database of the protein components of the mammalian stress granules. *Database (Oxford)* **2019,** (2019).
12. Millar, S. R., Huang, J. Q., Schreiber, K. J., Tsai, Y.-C., Won, J., Zhang, J., Moses, A. M. & Youn, J.-Y. A New Phase of Networking: The Molecular Composition and Regulatory Dynamics of Mammalian Stress Granules. *Chem Rev* (2023). doi:10.1021/acs.chemrev.2c00608
13. Zarnegar, B. J., Flynn, R. A., Shen, Y., Do, B. T., Chang, H. Y. & Khavari, P. A. irCLIP platform for efficient characterization of protein-RNA interactions. *Nat. Methods* **13,** 489–492 (2016).
14. Cox, J. & Mann, M. MaxQuant enables high peptide identification rates, individualized p.p.b.-range mass accuracies and proteome-wide protein quantification. *Nat. Biotechnol.* **26,** 1367–1372 (2008).
15. Feng, Z., Fang, P., Zheng, H. & Zhang, X. DEP2: an upgraded comprehensive analysis toolkit for quantitative proteomics data. *Bioinformatics* **39,** (2023).

16. Ou, J. & Zhu, L. J. trackViewer: a Bioconductor package for interactive and integrative visualization of multi-omics data. *Nat. Methods* **16,** 453–454 (2019).
17. Ge, X., Chen, Y. E., Song, D., McDermott, M., Woyshner, K., Manousopoulou, A., Wang, N., Li, W., Wang, L. D. & Li, J. J. Clipper: p-value-free FDR control on high-throughput data from two conditions. *Genome Biol.* **22,** 288–29 (2021).
18. Raudvere, U., Kolberg, L., Kuzmin, I., Arak, T., Adler, P., Peterson, H. & Vilo, J. g:Profiler: a web server for functional enrichment analysis and conversions of gene lists (2019 update). *Nucleic Acids Res.* **47,** W191–W198 (2019).
19. Ritchie, M. E., Phipson, B., Wu, D., Hu, Y., Law, C. W., Shi, W. & Smyth, G. K. limma powers differential expression analyses for RNA-sequencing and microarray studies. *Nucleic Acids Res.* **43,** e47–e47 (2015).
20. Kechin, A., Boyarskikh, U., Kel, A. & Filipenko, M. cutPrimers: A New Tool for Accurate Cutting of Primers from Reads of Targeted Next Generation Sequencing. *J Comput Biol* **24,** 1138–1143 (2017).
21. Smith, T., Heger, A. & Sudbery, I. UMI-tools: modeling sequencing errors in Unique Molecular Identifiers to improve quantification accuracy. *Genome Research* **27,** 491–499 (2017).
22. Sahadevan, S., Sekaran, T., Ashaf, N., Fritz, M., Hentze, M. W., Huber, W. & Schwarzl, T. htseq-clip: a toolset for the preprocessing of eCLIP/iCLIP datasets. *Bioinformatics* **39,** (2023).
23. Schwarzl, T., Sahadevan, S., Lang, B., Miladi, M., Backofen, R., Huber, W., Hentze, M. W. & Tartaglia, G. G. Improved discovery of RNA-binding protein binding sites in eCLIP data using DEWSeq. *Nucleic Acids Res.* **52,** e1 (2024).
24. Zhu, L. J., Gazin, C., Lawson, N. D., Pagès, H., Lin, S. M., Lapointe, D. S. & Green, M. R. ChIPpeakAnno: a Bioconductor package to annotate ChIP-seq and ChIP-chip data. *BMC Bioinformatics* **11,** 237–10 (2010).
25. Nystrom, S. L. & McKay, D. J. Memes: A motif analysis environment in R using tools from the MEME Suite. *PLoS Comput. Biol.* **17,** e1008991 (2021).
26. Gu, Z., Eils, R. & Schlesner, M. Complex heatmaps reveal patterns and correlations in multidimensional genomic data. *Bioinformatics* **32,** 2847–2849 (2016).
27. Love, M. I., Huber, W. & Anders, S. Moderated estimation of fold change and dispersion for RNA-seq data with DESeq2. *Genome Biol.* **15,** 550 (2014).
28. Gu, Z., Eils, R., Schlesner, M. & Ishaque, N. EnrichedHeatmap: an R/Bioconductor package for comprehensive visualization of genomic signal associations. *BMC Genomics* **19,** 234–7 (2018).
29. Robinson, J. T., Thorvaldsdóttir, H., Winckler, W., Guttman, M., Lander, E. S., Getz, G. & Mesirov, J. P. Integrative genomics viewer. *Nat. Biotechnol.* **29,** 24–26 (2011).
30. Abramson, J., Adler, J., Dunger, J., Evans, R., Green, T., Pritzel, A., Ronneberger, O., Willmore, L., Ballard, A. J., Bambrick, J., Bodenstein, S. W., Evans, D. A., Hung, C.-C., O'Neill, M., Reiman, D., Tunyasuvunakool, K., Wu, Z., Žemgulytė, A., Arvaniti, E., Beattie, C., Bertolli, O., Bridgland, A., Cherepanov, A., Congreve, M., Cowen-Rivers, A. I., Cowie, A., Figurnov, M., Fuchs, F. B., Gladman, H., Jain, R., Khan, Y. A., Low, C. M. R., Perlin, K., Potapenko, A., Savy, P., Singh, S., Stecula, A., Thillaisundaram, A., Tong, C., Yakneen, S., Zhong, E. D., Zielinski, M., Žídek, A., Bapst, V., Kohli, P., Jaderberg, M., Hassabis, D. & Jumper, J. M. Accurate structure prediction of biomolecular interactions with AlphaFold 3. *Nature* **630,** 493–500 (2024).

31. Narykov, O., Srinivasan, S. & Korkin, D. Computational protein modeling and the next viral pandemic. *Nat. Methods* **18,** 444–445 (2021).
32. Meng, E. C., Goddard, T. D., Pettersen, E. F., Couch, G. S., Pearson, Z. J., Morris, J. H. & Ferrin, T. E. UCSF ChimeraX: Tools for structure building and analysis. *Protein Sci* **32,** e4792 (2023).
33. Shen, S., Park, J. W., Lu, Z.-X., Lin, L., Henry, M. D., Wu, Y. N., Zhou, Q. & Xing, Y. rMATS: robust and flexible detection of differential alternative splicing from replicate RNA-Seq data. *Proc. Natl. Acad. Sci. U.S.A.* **111,** E5593–601 (2014).
34. Patro, R., Duggal, G., Love, M. I., Irizarry, R. A. & Kingsford, C. Salmon provides fast and bias-aware quantification of transcript expression. *Nat. Methods* **14,** 417–419 (2017).
35. Zhu, A., Srivastava, A., Ibrahim, J. G., Patro, R. & Love, M. I. Nonparametric expression analysis using inferential replicate counts. *Nucleic Acids Res.* **47,** e105 (2019).
36. Schindelin, J., Arganda-Carreras, I., Frise, E., Kaynig, V., Longair, M., Pietzsch, T., Preibisch, S., Rueden, C., Saalfeld, S., Schmid, B., Tinevez, J.-Y., White, D. J., Hartenstein, V., Eliceiri, K., Tomancak, P. & Cardona, A. Fiji: an open-source platform for biological-image analysis. *Nat. Methods* **9,** 676–682 (2012).
37. Perez-Riverol, Y., Csordas, A., Bai, J., Bernal-Llinares, M., Hewapathirana, S., Kundu, D. J., Inuganti, A., Griss, J., Mayer, G., Eisenacher, M., Pérez, E., Uszkoreit, J., Pfeuffer, J., Sachsenberg, T., Yilmaz, S., Tiwary, S., Cox, J., Audain, E., Walzer, M., Jarnuczak, A. F., Ternent, T., Brazma, A. & Vizcaíno, J. A. The PRIDE database and related tools and resources in 2019: improving support for quantification data. *Nucleic Acids Res.* **47,** D442–D450 (2019).
